# Global transcriptional regulation by cell-free supernatant of *Salmonella* Typhimurium peptide transporter mutant leads to inhibition of intra-species biofilm initiation

**DOI:** 10.1101/2020.07.15.204859

**Authors:** Kasturi Chandra, Prerana Muralidhara, Sathisha Kamanna, Utpal S. Tatu, Dipshikha Chakravortty

## Abstract

*Salmonella* is a genus of widely spread Gram negative, facultative anaerobic bacteria, which is known to cause ¼th of diarrheal morbidity and mortality globally. It causes typhoid fever and gastroenteritis by gaining access to the host gut through contaminated food and water. *Salmonella* utilizes its biofilm lifestyle to strongly resist antibiotics and persist in the host. Although biofilm removal or dispersal has been studied widely, the inhibition of the initiation of *Salmonella* biofilm remains elusive. This study was conducted to determine the anti-biofilm property of the cell-free supernatant obtained from a carbon-starvation inducible proline peptide transporter mutant (Δ*yjiY)* strain. Our study shows that *Salmonella* Δ*yjiY* culture supernatant primarily inhibits biofilm initiation by regulating biofilm-associated transcriptional network. This work demonstrates that highly abundant proteases such as HslV and GrpE cleave the protein aggregates, whereas global transcription regulators H-NS, FlgM regulate expression of SPIs and flagellar genes. Relatively low abundances of flavoredoxin, glutaredoxin, thiol peroxidase etc. leads to accumulation of ROS within the biofilm, and subsequent toxicity. This work further suggests that targeting these oxidative stress relieving proteins might be a good druggable choice to reduce *Salmonella* biofilm.

**Importance:** The enteric pathogen *Salmonella* forms biofilm in the internal organs of asymptomatic carriers, and on abiotic surfaces that leads to contamination of food and water. Biofilms are highly drug-resistant life forms that also helps in host immune evasion. Therefore, recent insurgence of drug tolerant strains necessitates development of biofilm inhibitory strategies, and finding novel druggable targets. In this study we investigated the bioactive molecules present in the cell-free supernatant of a biofilm deficient strain of *Salmonella* Typhimurium that inhibit biofilm initiation by the wildtype strain. Further we showed that the supernatant treatment leads to virulence defect *in vivo*. Collectively, our results suggest a comprehensive view of virulence regulation in *Salmonella* by the cell-free supernatant of the biofilm deficient strain.

## Introduction

*Salmonella* is one of the most common foodborne pathogens that causes one of the 4 major diarrheal diseases, salmonellosis (1, 2). The biofilm life form of this bacteria makes it a hardy human pathogen that can survive several weeks in a dry environment and many months in water (3). A biofilm refers to a community of bacteria adherent to a biotic/abiotic substratum, held together by its secreted extracellular polymeric substances. Biofilms are highly resistant to antibiotics, disinfectants etc. and biofilms on medical devices, catheters, implants are a major cause of hospital acquired infections (4, 5). Recent reports suggest that *Salmonella* can form biofilm on the gall bladder, leading to chronic infection (6, 7). The major constituents of *Salmonella* biofilm are curli, cellulose and BapA. Curli facilitates intercellular and cell to surface interactions, cellulose enables long range contact and provides characteristic stickiness to the biofilm and BapA strengthens curli interactions and stabilizes the biofilm (8). A major cue for biofilm formation is nutritional stress. Previously we found that the deletion of carbon starvation inducible gene, *yjiY*, leads to deficiency in biofilm formation (9). Antibiotic stress, DNA damage, cold stress, acid stress etc. can also regulate the transcription of *yjiY* gene (10–12). In *S*. Typhimurium, YjiY is known to regulate virulence by affecting flagellar class III genes, and the deletion of *yjiY* leads to reduced colonization in mice (9, 13), and upregulation of the virulence factor *mgtC*, leading to biofilm deficiency (14). This could indicate the possible role of YjiY in survival and defense against host induced stress in bacteria.

Recent studies corroborate that various culture supernatants of commensal bacteria can inhibit colonization and biofilm formation by their pathogenic counterparts. This inhibitory activity of culture supernatants can be seen across Gram positive, and Gram negative bacteria as well as across kingdoms. Iwase *et al*. showed that *Staphylococcus epidermidis* Esp inhibits biofilm formation and nasal colonization by pathogenic *Staphylococcus aureus* (15). In another study, probiotic *Escherichia coli* Nissle (EcN) inhibited biofilm formation in pathogenic EHEC, by a secreted bifunctional (protease and chaperone) protein DegP (16). There are various other modes of interspecies biofilm regulation such as biofilm dispersal proteins (17), indole-mediated biofilm regulation (18), quorum sensing molecules, and antimicrobial peptides (19). The inhibition can also be interkingdom, such as *Aspergillus* biofilm inhibition by *Pseudomonas* culture supernatants (20). While there are a few strategies to deal with preformed biofilms (21, 22), the inhibition of biofilm initiation is comparatively less explored in human pathogens. In our lab we found that *Salmonella* WT could not form biofilm when cocultured with Δ*yjiY* strain. This study was designed to investigate the novel biofilm inhibitory activity of biofilm deficient strain *Salmonella* Typhimurium Δ*yjiY*.

In this study, we are reporting a novel mechanism of biofilm initiation inhibition by a complex transcriptional regulatory network. This complex network causes adhesion impairment, flagellar motility inhibition, cleavage of protein aggregates and quorum sensing inhibition. Furthermore, we have shown that STM Δ*yjiY* supernatant can also inhibit biofilm formation by *E. coli*. We have also shown that the supernatant treatment can impair the invasion of the pathogen in the *C. elegans* gut, thus reducing its virulence.

## Results

### *Salmonella ΔyjiY* culture supernatant inhibits WT biofilm formation

To understand the biofilm inhibitory property of STM Δ*yjiY* culture supernatant, we inoculated WT bacteria in biofilm inducing low salinity media with or without the culture supernatant of different strains. The Δ*yjiY* bacterial culture supernatant exhibited the biofilm inhibitory property (**Fig. 1A**), which was absent in the supernatant of the strain where *yjiY* was trans-complemented in a plasmid (STM Δ*yjiY:pQE60-yjiY)*. Previous study showed that Δ*yjiY* is a biofilm deficient strain (14). To check whether the biofilm inhibitory property is specific to Δ*yjiY* supernatant, we inoculated WT strain with biofilm deficient strain Δ*csgD* (**Fig. S1A**). We observed inhibition only in the case of Δ*yjiY* supernatant indicating that the inhibitory molecules are unique to the Δ*yjiY* secretome, and are independent of the activity of CsgD. To further validate that the inhibition is a cell-free phenomenon, we inoculated WT bacteria along with either live cells of other strains along with spent culture media (coculture), supernatant-free cell pellets, cell free supernatant, or whole cell lysate, and quantified the biofilm on solid-liquid-air interface. Although Δ*yjiY* showed significant inhibition of biofilm formation in all the setups, the maximum inhibition was observed with Δ*yjiY* culture supernatant (**Fig. 1B**). The supernatants were concentrated using an Amicon ultra filter device and total protein was quantified. We found that minimum biofilm inhibitory concentration (**MBIC**; minimum concentration of total protein required to inhibit biofilm formation by WT strain) falls between 15-20 ng protein/ml (**Fig. 1C**), therefore we used supernatant containing 20 ng protein/ml of supernatant for further experiments. We did not find any difference in the growth (**Fig. S1B**), suggesting that the supernatant lacks any bactericidal or bacteriostatic properties. Interestingly, we observed slower, yet longer exponential growth after supernatant treatment (**Fig. S1C**). To determine whether the secretion of the inhibitory component(s), is dependent on the culture media, we grew the bacteria in minimal media (M9 media supplemented with 0.5% glucose) that exerts nutrition stress and used the supernatant to treat WT bacteria. We observed that the biofilm inhibitory molecule(s) were active even in minimal media (**Fig. 1D**), suggesting that the production of the inhibitory molecule(s) is not dependent on the nutritional condition and is an intrinsic property of the Δ*yjiY* strain. Additionally we checked the temporal expression or accumulation of the inhibitory component(s) by inoculating the WT strain with culture supernatant harvested from 2, 3, 4 and 5 day old Δ*yjiY* cultures. Our results suggest that the optimum concentration of the inhibitory component(s) is/are reached after 3 days of growth (**Fig. S1D**), which remains unchanged on the 4^th^-5^th^ days of growth. Therefore, we used filtered culture supernatant from 3 day old Δ*yjiY* culture for further experiments.

**Fig 1.**
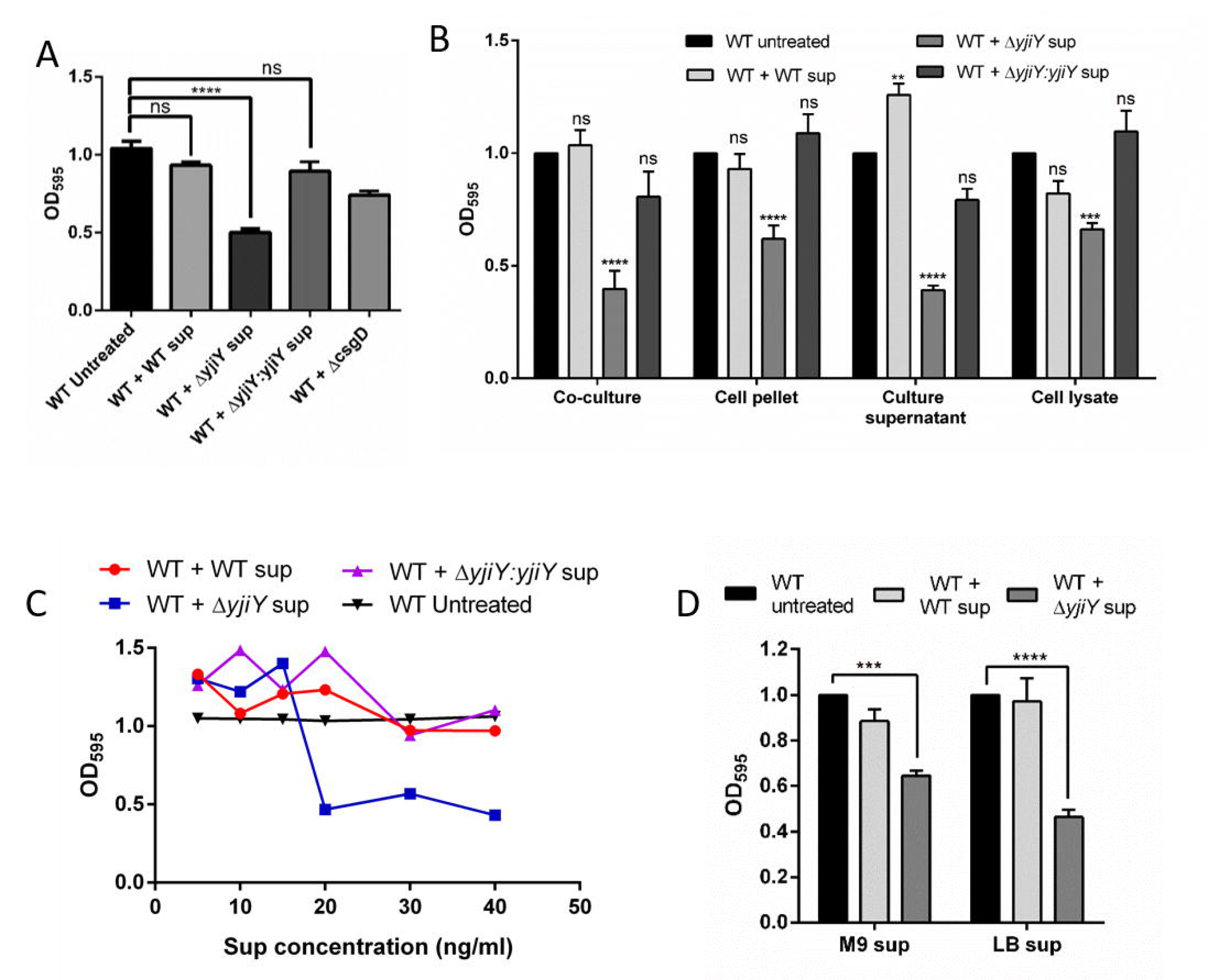
*Salmonella* biofilm deficient Δ*yjiY* inhibits biofilm formation by WT strain. A. Biofilm formation ability of the WT strain after 72 hours of inoculation in presence or absence of WT, Δ*yjiY, ΔyjiY:yjiY* and Δ*csgD* culture supernatants was checked. Crystal violet staining was carried out to check the biofilm formed on the solid-liquid interphase (Data are presented as mean + SEM of 12 independent experiments). B. Similarly biofilm formation assay was carried out in presence or absence of overnight grown different bacterial strains (co-culture), washed bacterial pellet (cell pellet), cell free supernatant (culture supernatant) and cell lysates (Data are presented as mean + SEM of 4 independent experiments). C. Different concentration of the cell free supernatant was used to check the minimum biofilm inhibitory concentration (MBIC) of Δ*yjiY* supernatant (Data are presented as mean + SEM of 4 independent experiments). D. Biofilm inhibitory property of the Δ*yjiY* supernatant was checked with supernatant isolated from bacteria grown in shaking condition in LB media and minimal M9 media (Data are presented as mean + SEM of 4 independent experiments). Student’s t-test was used to analyze the data; p values ****<0.0001, ***<0.001, **<0.01, *<0.05.

### Δ*yjiY* culture supernatant weakens the WT biofilm and interferes with cell structure

The characteristic EPS components are cellulose (produced by *bcsA* encoded cellulose synthase), curli fimbriae (encoded by *csgAB)*, BapA and LPS (23). We observed a significant reduction of the EPS bound Congo red fluorescence intensity (**Fig. 2A, S2A**) and biofilm thickness (**Fig. S2B, S2C**) upon Δ*yjiY* supernatant treatment. We also found that Δ*yjiY* supernatant treatment significantly reduced the strength of the biofilm pellicle (**Fig. S2D, S2E, 2B**) than that of untreated or WT supernatant treated samples. SEM and AFM analysis of the biofilm surface showed that the Δ*yjiY* supernatant treated biofilms lack the characteristic dome shape of a proper biofilm (**Fig. 2C, S2F**). We also noticed that the median cell length increased upon Δ*yjiY* supernatant treatment (1.58+0.30 μm) as compared to the untreated WT cells (1.38+0.34 μm) (**Fig. 2D, 2E, S2G**), hinting towards the presence of multiple regulatory components in the Δ*yjiY* supernatant that can modulate multiple phenotypic effects.

**Fig 2.**
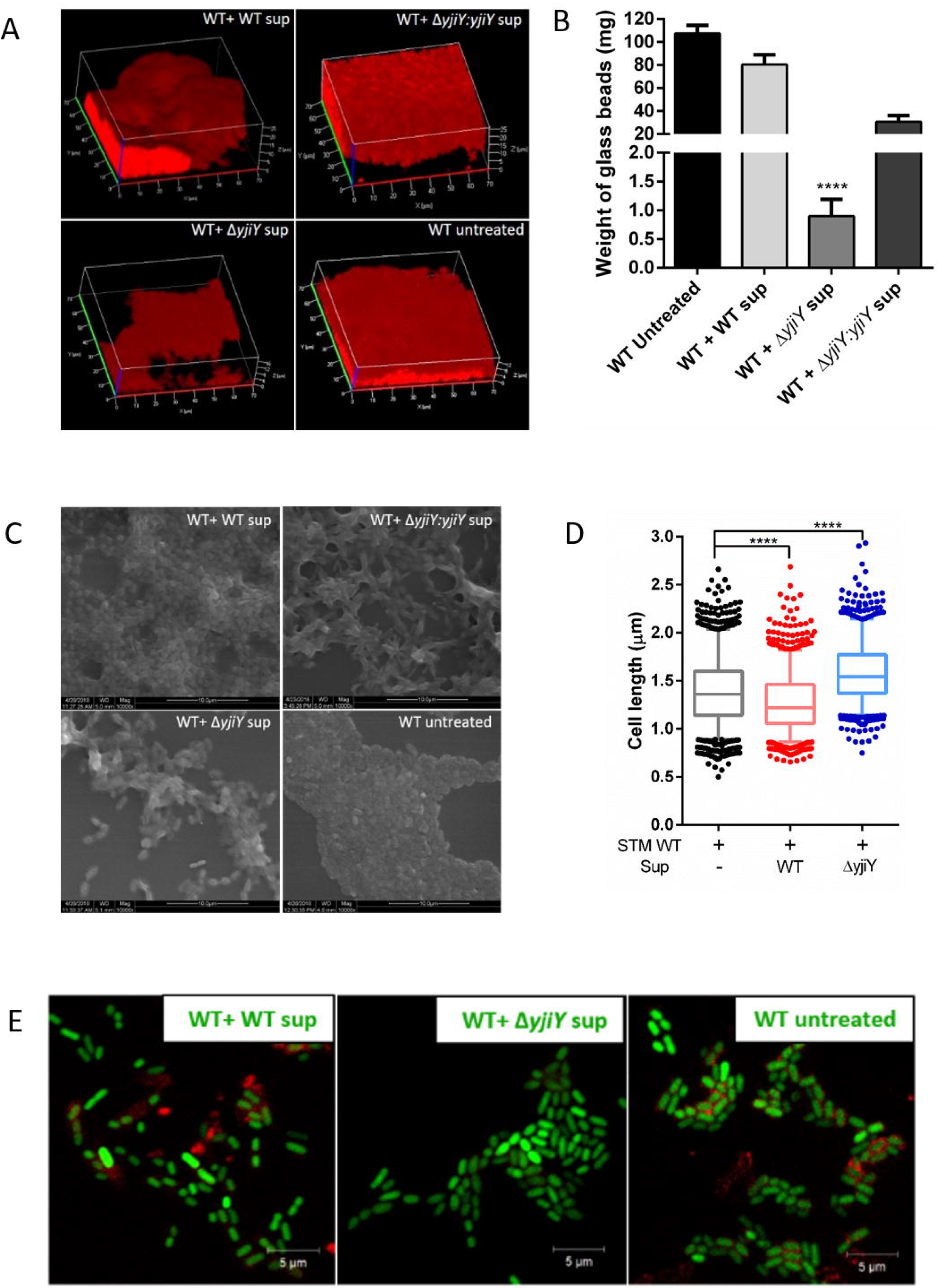
*Salmonella ΔyjiY* cell free supernatant significantly reduces the biofilm biomass by reducing cell-cell adhesion. A. Representative images of biofilm formed on coverslips that were stained with Congo red and imaged using a confocal microscope to generate 3D images and quantify cellulose biomass. Scale is shown on the X- and Y-axes. B. The tensile strength of the biofilm was measured by glass bead assay. Weight of glass beads required to just sink the biofilm to bottom, was plotted (Data are presented as mean + SEM of 5 independent experiments). C. Representative scanning electron micrograph of biofilm formed on a coverslip. Scale bar is 10 μm. D. Cell length of the biofilm inoculated treated or untreated STM WT cells was measured using ImageJ, and plotted (Data are presented as mean + SEM of 1200 cells were measured from 3 independent experiments). E. Representative confocal images of biofilm cells showing a difference in cell length. Scale bar is 5 μm (1000-1200 cells were measured from 3 independent experiments for each treatment). Student’s t-test was used to analyze the data; p values ****<0.0001, ***<0.001, **<0.01, *<0.05.

### The active molecule(s) is/are protein(s)

Previous studies have shown that secreted components from some bacteria can inhibit biofilm formation by the wild type strain or closely related species (15, 16, 20). To delineate the chemical nature of the inhibitory molecule(s) present in Δ*yjiY* supernatant, we treated the supernatant with chemical agents, such as a divalent cation chelator (EDTA), protease (Proteinase K), protease inhibitor (PMSF), RNase and DNase and quantified the biofilm inhibition. We found that upon pre-treatment with different concentrations of EDTA, the inhibitory property remained intact. Interestingly, 10 mM EDTA enhanced biofilm formation with both treated and untreated WT strain (**Fig. 3A**). Since EDTA is known to chelate divalent cations and inhibit a few proteases at higher concentrations (24–27), our data signify the requirement of divalent cations and/or active proteases for biofilm inhibition. Upon treating the supernatants with 20 mg/ml Proteinase K, the ability of Δ*yjiY* supernatant to inhibit biofilm formation was significantly reduced, suggesting that the inhibitory molecule(s) are protein(s) (**Fig. 3B**). Since the activity of many proteins is sensitive to even small changes in pH and temperature, we checked the activity of Δ*yjiY* supernatant at different pH and temperatures. Surprisingly, we found that the active component(s) is/are heat stable at 65 °C and 95 °C temperatures (**Fig. 3C**) and stable over a wide range of acidic and alkaline pH (**Fig. 3D**), although there was a small reduction in biofilm inhibition at pH 9.0. As recent studies show that several small noncoding RNAs regulate biofilm formation and other virulence traits in *Vibrio cholerae* and *Pseudomonas aeruginosa* (28, 29), we tested the stability of the component(s) after treating the supernatant with RNase. Although the inhibition was lost upon RNase treatment and incubation at 37°C, we found a similar loss of inhibition with only heating the supernatant at 37°C (**Fig. S3A**), suggestive of the heat, rather RNase treatment, to be the reason for the loss of inhibition. We also fractionated the supernatant using an Amicon 3k MWCO ultra filter device, and we found that the active component(s) of Δ*yjiY* supernatant is/are >3kDa molecular weight (**Fig. S3B**), quashing the role of small molecules and ions in biofilm inhibition by Δ*yjiY* supernatant. Since the inhibitory activity was abolished upon proteinase treatment, we attributed the inhibition to protein components and quantified the total protein for further experiments.

**Fig 3.**
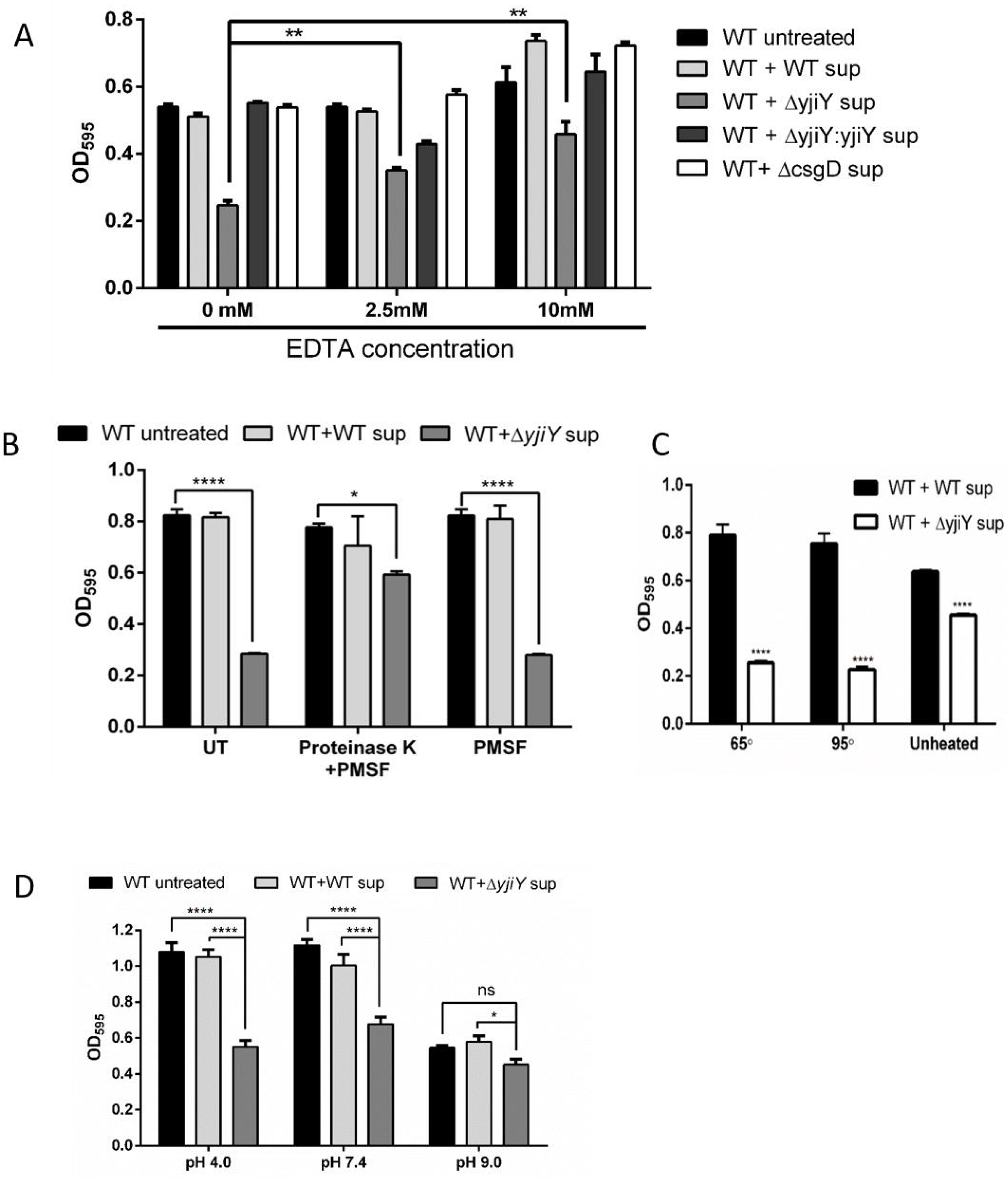
The physicochemical property of the inhibitory molecules is proteinaceous. A. The cell-free supernatants were treated with different concentrations of EDTA and checked for their ability to inhibit biofilm formation by STM WT (Data are presented as mean + SEM of 3 independent experiments). B. The supernatants were treated with proteinase K, and heated to 37 °C for 1 hour, and checked for the biofilm inhibitory activity. PMSF was used to inactivate proteinase K after 1 hour, and was also used as a control (Data are presented as mean + SEM of 3 independent experiments). C. The supernatants were heated to 65 °C and 95 °C to check the thermostability of the active component(s), followed by biofilm inoculation with the treated or untreated supernatants (Data are presented as mean + SEM of 3 independent experiments). D. The pH sensitivity of biofilm inhibitory action of the supernatants were checked. The pH of the supernatants as well as the biofilm media was made acidic (pH 4.0) or alkaline (pH 9.0) with concentrated HCl or NaOH, respectively, and checked for the activity of the active molecule(s) (Data are presented as mean + SEM of 3 independent experiments). Two-way ANOVA and Student’s t-test were used to analyze the data; p values ****<0.0001, ***<0.001, **<0.01, *<0.05.

### Active molecules inhibit biofilm only during the initial phases, and cannot disrupt mature biofilm

To determine the effect of the active molecule on mature biofilm, we treated mature biofilm pellicles with the supernatants and checked for dispersion. Our results showed that the Δ*yjiY* supernatant could not disrupt pre-formed biofilm, hinting towards the effect of the active molecule(s) on biofilm initiation (**Fig. 4A**). To determine whether the supernatant treated WT cells remained biofilm defective in the absence of the inhibitory molecules, the Δ*yjiY* supernatant treated WT cells were re-inoculated in biofilm medium without the supernatant, and monitored for pellicle formation. We observed that the inhibition was diminished in the absence of the supernatant (**Fig. 4B**), although the defect reappeared when these cells were re-treated with the inhibitory supernatant. Together our data suggest that supernatant mediated initial molecular reprogramming is required for biofilm inhibition, and that inhibitory molecules do not alter the inherent biofilm forming ability of the WT cells.

**Fig 4.**
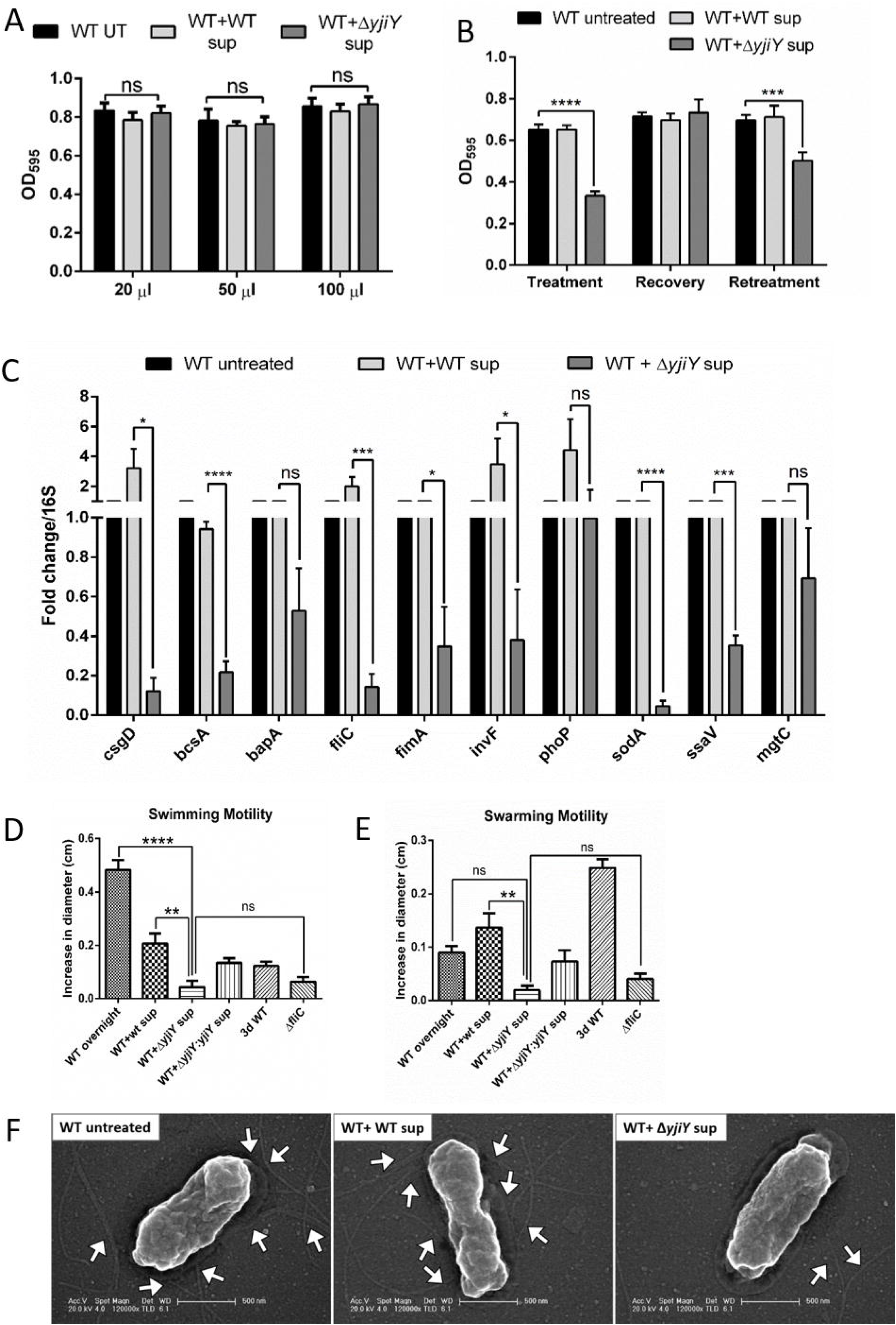
The supernatant interrupts biofilm formation by transcriptional switching during initiation of micro-colony formation. A. Varied concentrations of the STM Δ*yjiY* supernatant were used to pre-formed mature biofilm of STM WT and the ability of the supernatant to disintegrate the biofilm was checked (Data are presented as mean + SEM of 5 independent experiments). B. The extent of inhibitory effect of the supernatants were checked by recovery and repeated exposure of the WT strain to the supernatant (Data are presented as mean + SEM of 3 independent experiments). C. RNA was isolated from untreated STM WT cells or WT cells treated with the indicated supernatants for 72 hours. RTPCR was done for indicated genes, and C_T_ values were first normalized to 16s rRNA gene and then normalized to that of untreated WT cells. Y axis values above and below 1 denotes upregulation and downregulation, respectively (Data are presented as mean + SEM of 3 independent experiments). (D, E). *In vitro* motility of the treated and untreated STM WT cells were checked on 0.3% swim agar plates or 0.5% swarm agar plates. The radius of the motile zone was imaged after different time, measured using ImageJ and plotted (Data are presented as mean + SEM of 4 independent experiments). F. The flagellar status of the treated or untreated cells (approximately 50 cells in each samples, in two different experiments) was imaged using scanning electron microscope. White arrows show the flagella. Scale bar is 500 nm. Two-way ANOVA and Student’s t-test were used to analyze the data; p values ****<0.0001, ***<0.001, **<0.01, *<0.05.

### The inhibitory molecules impair flagella-mediated initial attachment to abiotic surfaces

*Salmonella* biofilm formation can be divided into 4 distinct steps, (i) initial attachment to the abiotic surface, (ii) secretion of adhesive components, (iii) maturation and (iv) dispersion of biofilm upon relief of the stress (30). Since biofilm initiation was inhibited and retained beyond 72 hours, we checked the expression of biofilm-associated genes (*csgD, bcsA, fliC)*, SPI-1 effectors (*invF, sopD*) and virulence factors (*phoP, sodA, mgtC*) from untreated or treated WT cells, 72 hours post inoculation. We found that *csgD, bcsA, fliC* and *invF* expression was significantly downregulated in Δ*yjiY* supernatant treated WT cells (**Fig. 4C**). Since, flagella-mediated initial attachment initiates biofilm formation (31, 32), we investigated the effect of Δ*yjiY* supernatant on flagellar motility. We observed that Δ*yjiY* supernatant treatment significantly reduced both swimming and swarming motility (**Fig. 4D, 4E** and **S4A**), indicating a flagella-mediated motility defect. We further observed the downregulation of *fliC* occurs as early as 4-6 hours post inoculation with the Δ*yjiY* supernatant (**Fig. S4B**). Interestingly, we observed an increase in *fliC* expression at 12 hours post inoculation. We reasoned that adhesion deficiency arises from the sequential effect of the initial downregulation of*fliC*, followed by an increase in the planktonic population. SEM analysis showed that Δ*yjiY* supernatant treated bacteria had significantly fewer flagella (**Fig. 4F, S4C**). Altogether, these results indicate a defect in flagella-mediated initial attachment of the bacteria, sets the course for biofilm deficiency.

### Δ*yjiY* supernatant reduces adhesion and virulence of *Salmonella* both *in vitro* and in *C. elegans* gut

Initial attachment of *Salmonella* on the host gut epithelial cells requires swimming through the mucus layer (33). SPI1 encoded Invs, Sops and Sips are important in initial invasion (34). Since Δ*yjiY* supernatant treatment downregulated *invF* and *sopD* expression, we checked the infectivity of the supernatant treated cells in mammalian intestinal epithelial cells, Int407. Invasion assay shows that Δ*yjiY* supernatant treated cells are defective in initial invasion, although these cells showed significantly higher intracellular proliferation than the untreated cells (**Fig. 5A** and **5B**), suggesting that the Δ*yjiY* supernatant only makes WT cells invasion defective by inhibiting flagella-mediated adhesion. To check the infectivity of the Δ*yjiY* supernatant treated cells in a systemic condition, we fed young adult *C. elegans* N2 worms RFP-STM-WT bacteria and quantified gut colonization. The micrograph images and CFU analysis show that Δ*yjiY* supernatant treated RFP-STM-WT cells were able to colonize the gut lumen when fed continuously, but did not persist (**Fig. 5C, 5D, 5E**). Therefore, our data suggest that Δ*yjiY* supernatant treatment impairs *in vitro* invasion and *in vivo* colonization.

**Fig 5.**
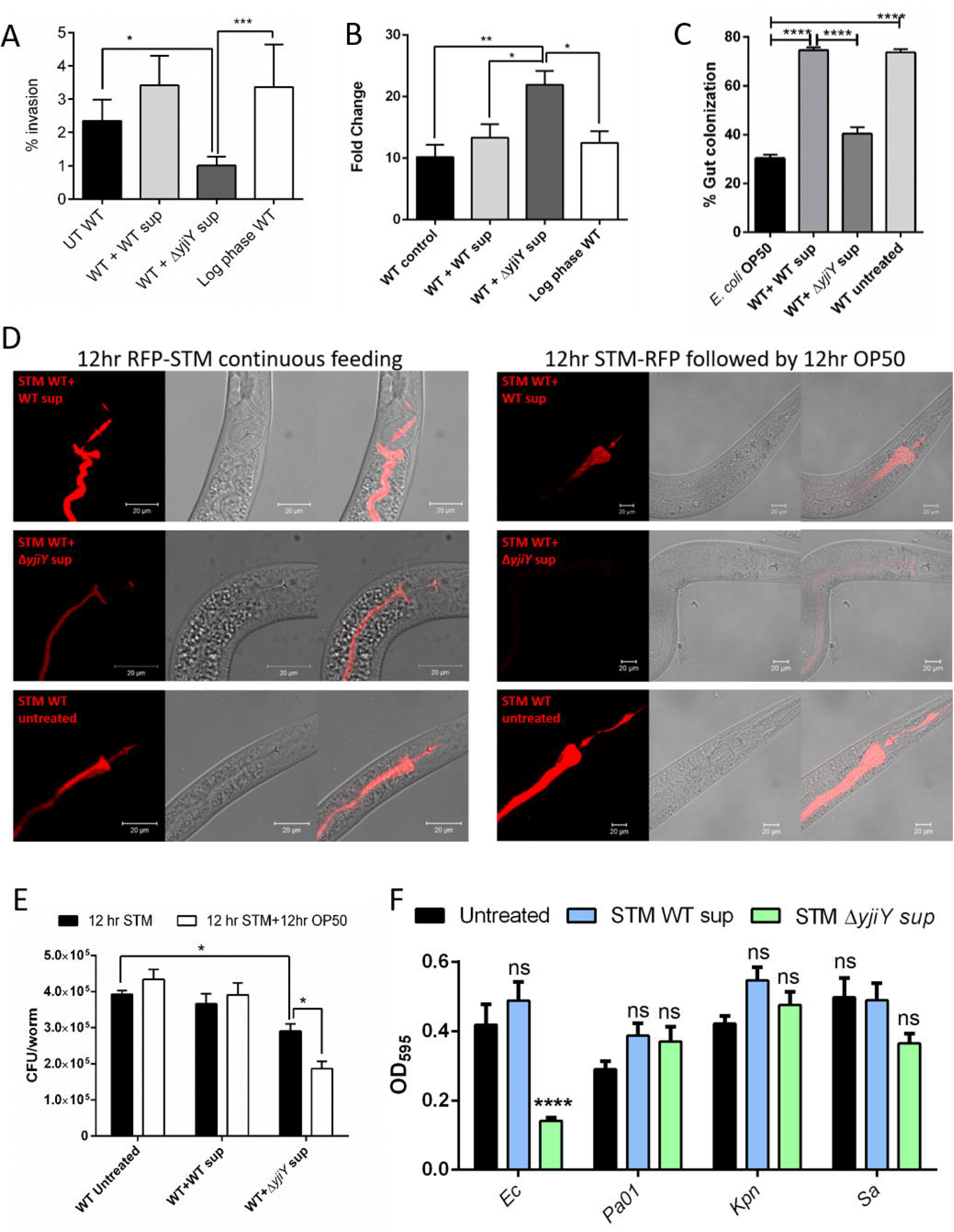
STM Δ*yjiY* supernatant treatment impairs virulence of STM WT. A. *In vitro* invasiveness of the supernatant treated WT bacteria was checked by infecting adherent human epithelial Int407 cell line and percent invasion was plotted. Log phase WT culture was used as a control (Data are presented as mean + SEM of 3 independent experiments). B. Survival of intracellular *Salmonella* in Int407 cells was checked by gentamicin protection assay. The intracellular bacterial fold proliferation was measured as the ratio of intracellular bacterial CFU after 16 hours to the CFU at 2 hours (Data are presented as mean + SEM of 3 independent experiments). C. Percent gut colonization was measured by measuring the diameters of the colonized gut and total body width after feeding the worms continuously for 12 hours with *mCherry-E.coli* OP50 or supernatant treated/untreated mCherry-STM WT (approximately total 25 worms were measured in 2 independent experiments). D. Representative images of 3 days supernatant treated bacterial colonization in young adult *C. elegans* N2 worms. Bacterial colonization in the gut lumen was checked after 12 hours of continuous STM WT feeding and/or 12 hours STM WT feeding, followed by 12 hours *E. coli* OP50 feeding (approximately total 25 worms were imaged in 2 independent experiments). Scale bar is 20 μm. E. Bacterial CFU from infected *C. elegans* was determined by lysing 10 worms (in triplicates) in M9 buffer followed by dilution plating (Data are presented as mean + SEM of 2 independent experiments). Student’s t-test was used to analyze the data; p value *<0.05. F. Biofilm inhibitory effect of STM Δ*yjiY* supernatant on few other enterobacteriaceae pathogens was checked by inoculation the strains with 1% v/v supernatants (Ec-*Escherichia coli* DH5α, Pa01-*Pseudomonas aeruginosa* strain PA01, Kpn-*Klebsiella pneumoniae*, Sa-*Staphylococcus aureus*, Data are presented as mean + - SEM of 3 independent experiments). Two-way ANOVA and Student’s t-test were used to analyze the data; p values ****<0.0001, ***<0.001, **<0.01, *<0.05.

### Inhibitory molecules impede biofilm formation in closely related species

We next checked the effectiveness of STM Δ*yjiY* supernatant on the biofilms of common human pathogens. We inoculated *E. coli* DH5α, *Pseudomonas aeruginosa* PA01, *Klebsiella pneumoniae, and Staphylococcus aureus* wild-type strains with STM WT supernatant, and STM Δ*yjiY* supernatant, and quantified the biofilm. We found that the STM Δ*yjiY* supernatant significantly inhibited only *E. coli* DH5α biofilm which is regulated by CsgD (**Fig. 5F**). Therefore, we concluded that the Δ*yjiY* supernatant can cross-react with closely related Enterobacteriaceae, *E. coli*, but is not effective against distant members such as *Klebsiella pneumoniae* or different family (*Pseudomonas aeruginosa*) or phylum (*Staphylococcus aureus)*.

### The active components are primarily global transcription factors that regulate multiple cellular processes

To further identify the biofilm inhibitory molecule(s), concentrated supernatants from WT and Δ*yjiY* cultures were resolved on 10% SDS-PAGE. After colloidal CBB staining, we clearly visualized 14 differential bands in Δ*yjiY* supernatant compared to WT supernatant (**Fig. 6A**). We next analyzed the secretome in LC Q-TOF MS/MS. Among the 244 proteins, 188 proteins were present in both supernatants at differential levels, while 38 and 58 proteins were enriched in the WT supernatant and Δ*yjiY* supernatant, respectively (**Fig. 6B**). These proteins may have resulted from cell lysis during growth or they were secreted in the supernatant via an active secretion system. After careful data-mining, among the proteins found only in Δ*yjiY* supernatant (**Table S1**), probable transcriptional regulatory protein YebC, anti-sigma28 factor FlgM, a serine protease inhibitor ecotin and transcription termination/anti-termination protein NusG were selected for further analysis. Many transcriptional regulators (H-NS, Rnk, StpA), cold shock proteins (CspE, CspC), chaperones GrpE, ATP-dependent protease HslV, proteins related to oxidative stress and iron homeostasis (FldA, SodB, YdhD, Tph and Ftn, Bcp, Bfr, respectively) were differentially present in both supernatants (**Table S2**). Additionally, we treated STM WT cells with purified ecotin, HNS, and NusG (**Fig. S5**), to delineate the effect of each of these proteins. Interestingly, the fractions containing monomeric ecotin and NusG, as well as homodimeric HNS (**Fig. S6A, S6B**), showed biofilm inhibition similar to that of Δ*yjiY* supernatant, whereas the buffer (50mM Tris) treated WT cells did not have biofilm defect (**Fig. 6C**). Together, our data suggest that Δ*yjiY* supernatant perturbs various cellular processes leading to biofilm defect by complex transcriptional regulation (**Fig. 6D**).

**Fig 6.**
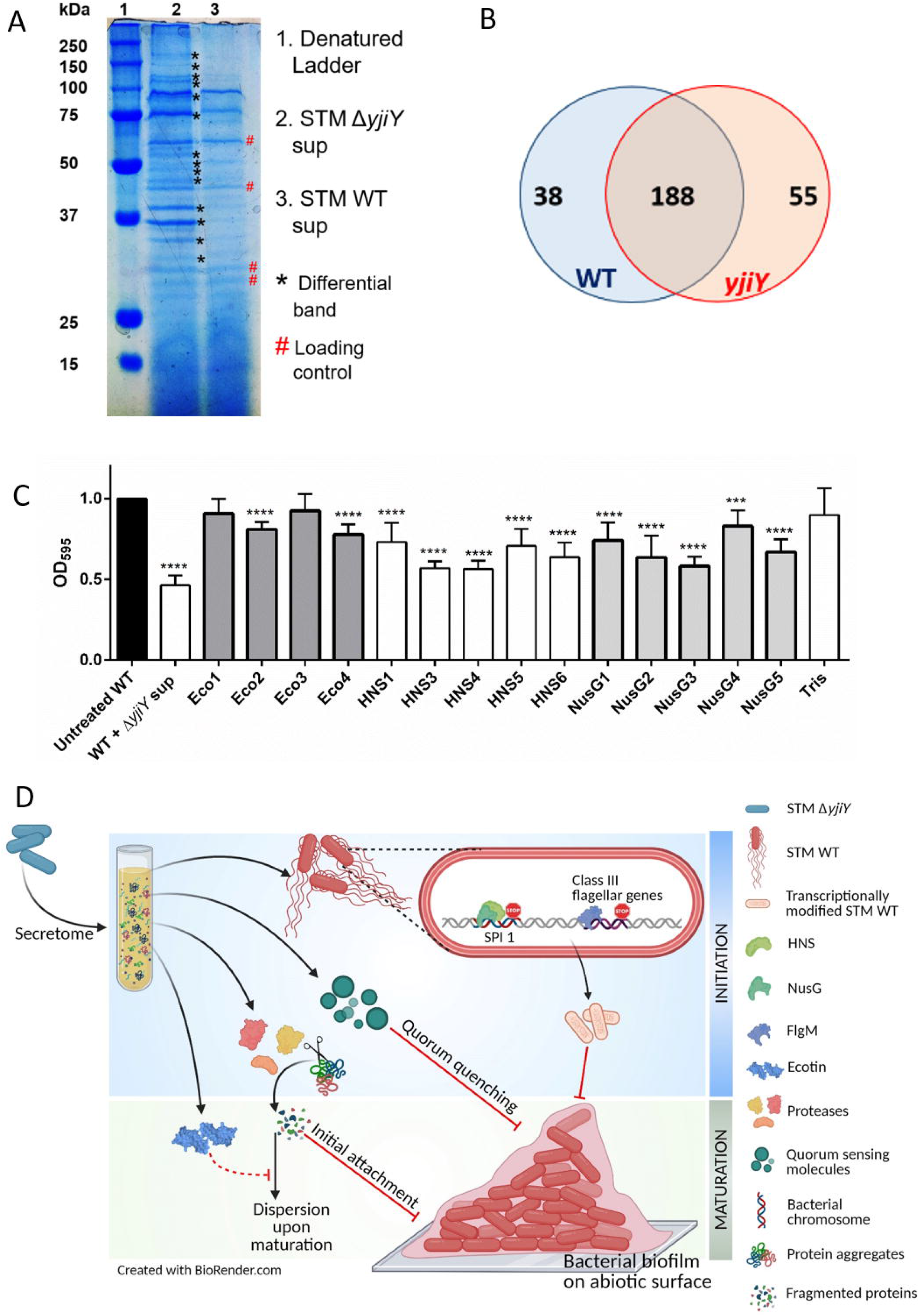
Global transcriptional regulator HNS along with NusG causes major transcriptional regulation leading to biofilm inhibition and motility inhibition. A. SDS PAGE of the STM WT and STM Δ*yjiY* supernatant showing presence of multiple differential bands in the supernatants. B. Venn diagram showing proteins present in the supernatants detected in LC QTOF MS/MS analysis, blue and red circles represent number of proteins found in STM WT and Δ*yjiY* culture supernatants, respectively. C. Biofilm formation ability of the WT strain after 72 hours of inoculation in presence of Δ*yjiY* culture supernatant and 200ng ecotin/HNS/NusG fractions was checked. Crystal violet staining was carried out to check the biofilm formed on the solid-liquid interphase (Data are presented as mean + SEM of 3 independent experiments; One-way was used to analyze the data; p values ****<0.0001, ***<0.001). D. Proposed model of biofilm inhibition by Δ*yjiY* supernatant. The model depicts that transcription factors, proteases, QS-quenching molecules, and protease inhibitors are secreted actively or released during cell lysis in Δ*yjiY* supernatant, which upon entering into the WT cells by either active transport or membrane diffusion, regulates biofilm formation and motility.

## Discussion

*Salmonella* forms biofilm to evade host defense, colonize in host, persist in asymptomatic host, and transmit to a new host (35). The transition from planktonic to biofilm mode depends upon various stress factors. The master regulator CsgD controls the production of EPS components such as cellulose, tafi (curli fimbriae) etc. by regulating AdrA. BapA is important both for biofilm production and attachment to intestinal epithelium (36). In this study we have shown how metabolic stress determines the fate of the infectious WT strain. From preliminary data, we found that coculture of STM WT and Δ*yjiY* led to biofilm defect in the WT strain. Although various studies have shown that interspecies and intraspecies biofilm inhibitions exist in nature (15, 20, 37), in *Salmonella*, this phenotype is novel. Therefore, we carried out experiments to understand the underlying mechanism.

In this study, we found that biofilm inhibition was unique to Δ*yjiY* supernatant and the inhibition was reversed when WT cells were treated with culture supernatant from the complement strain (STM Δ*yjiY:pQE60-yjiY)*. Although we observed that the Δ*yjiY* supernatant had only a biofilm inhibitory effect and lacked bactericidal properties (as observed from the growth analysis), STM WT cells showed delayed yet prolonged exponential growth upon supernatant treatment. Furthermore, the exopolysaccharide cellulose was sparse in the biofilm of Δ*yjiY* supernatant treated WT cells. Since only cellulose mutant *Salmonella* Typhimurium was proficient in adhering to tumour cells (38), our data imply the presence of multiple inhibitory factors involved in the inhibition by the supernatant. While incubation of Δ*yjiY* supernatant at 37°C was found to reduce the inhibition, heating at 65 °C and 95 °C enhanced the inhibition, indicating that the inhibitory molecules might be heat-shock proteins or high temperature inducible stress proteins. Fascinatingly we observed that the effect of the inhibitory molecules is temporary, and they do not cause any genetic change to alter the inherent biofilm forming competency of the WT cells. Δ*yjiY* supernatant was also found to exert its anti-biofilm activity only when administered at the beginning of biofilm formation, suggesting a modification in the complex network through which biofilm develops. In the Δ*yjiY* supernatant treated WT cells, the expression of biofilm genes and virulence genes was reduced significantly. *sodA* downregulation suggests that these cells are prone to ROS assault. A previous study from our group showed that YjiY depletion upregulates *mgtC* leading to biofilm defect (14). Interestingly, we observed downregulation of *mgtC*, implying that biofilm inhibition by Δ*yjiY* supernatant and the inherent biofilm defect of the Δ*yjiY* strain follow different mechanisms. The Δ*yjiY* supernatant treated WT cells showed significantly less invasion in Int407 cells. It was recently proposed that *Salmonella* persists in *C. elegans* gut by forming biofilm (39). Δ*yjiY* supernatant treated WT cells showed a concomitant reduction in colonization in the worm gut, suggesting an *in vivo* biofilm defect. We also observed the absence of flagella and other protein aggregates upon Δ*yjiY* supernatant treatment, leading to defects in cell aggregate formation and biofilm initiation. While there was a temporal increase in *fliC* in Δ*yjiY* supernatant treated WT cells, these cells exhibited defective motility after 72 hours of treatment, further validating the importance of flagella-mediated motility in the initial attachment of the bacteria to the substratum. Interestingly, we found that STM Δ*yjiY* supernatant effectively inhibited *E. coli* biofilm formation. Although *Salmonella* diverged from *E. coli* by acquiring virulence-associated genes, they share many evolutionarily conserved cellular pathways (40–42). Therefore this observation reiterates that the inhibitory effects are more profound among closely related species. In the proteomics analysis of the Δ*yjiY* supernatant, we specifically detected three potential inhibitory candidates: proteinase K sensitive ecotin (43), anti-sigma28 factor FlgM (a negative transcriptional regulator of class III flagellar genes (44, 45)) and YebC (negatively regulates quorum sensing in *Pseudomonas aeruginosa* PA01 (46)). The abundance of FlgM correlates with the absence of flagella in the Δ*yjiY* supernatant treated WT cells. NusG works synergistically with the global transcriptional regulator H-NS, which binds specifically to AT-rich SPIs in the *Salmonella* genome and represses those genes (47, 48). Interestingly, in *E. coli*, H-NS has been linked to the cell cycle, since the cells attempt to optimize spatial H-NS concentration by maintaining a constant ratio of H-NS to chromosomal DNA in the cell (49), which might explain the increase in cell length after Δ*yjiY* supernatant treatment. HslV, a heat-inducible ATP-dependent protease subunit of a proteasome-like degradation complex (50) and GrpE, which is involved in removal of protein aggregates leading to unsuccessful biofilm formation (51) were found in higher abundance in Δ*yjiY* supernatant. Doyle *et al*.(52) showed that excess GrpE inhibits the interaction between DnaK and the regulatory protein, RepA. Since RepA helps maintain the plasmid copy number in *E. coli* (53), inactivation of RepA might explain cell elongation upon Δ*yjiY* supernatant treatment. While protein aggregation is necessary during initial attachment of the bacteria to an abiotic surface, upon maturation, proteolytic activity of several proteases helps in dispersal of biofilm (54). Presence of a protease inhibitor, such as ecotin, might hinder this process, leading to dysregulation of biofilm maturation and dispersion. Although several moonlighting functions have been demonstrated for mycobacterial superoxide dismutase and DnaK in modulating host response (55), any such activity for other hits remains elusive.

Oxidative stress proteins such as flavoredoxin, SodB, glutaredoxin, thiol peroxidase (Tph), and thioredoxin-dependent thiol peroxidase (Bcp) were less abundant in Δ*yjiY* supernatant, among which Tph and SodB help reduce oxidative stress in STEC biofilm (56). In *E. faecalis*, Tph is required for the *in vitro* oxidative stress response and survival inside murine macrophages (57). Iron availability has been shown to both positively and negatively regulate biofilm formation through complex network systems (58). Although high Fe can lead to ROS production, Kang and Kirienko suggested that iron uptake and homeostasis are essential for successful biofilm formation in *Pseudomonas aeruginosa* (59). The Fe-storage proteins bacterioferritin (Bfr) and ferritin A (Ftn), required to prevent ROS generation via the Fenton reaction and DNA damage, are functionally very large proteins with a core to accommodate 3000 Fe atoms. Fe-rich conditions induce *E. coli* FtnA and Bfr due to loss of repression by small RNA RyhB(60). Similarly, *Salmonella* Typhimurium Bfr is involved in reducing intracellular Fe toxicity, and the absence of Bfr causes increased intracellular free Fe^2+^ ion and oxidative stress(61).

Therefore, we conclude that Δ*yjiY* supernatant treatment inhibits biofilm formation by WT in four major ways- (i) NusG-HNS mediated transcriptional repression of AT-rich SPI-encoded genes, making the WT bacteria less virulent; (ii) FlgM mediated downregulation of class III flagellar genes, impairing flagella-mediated initial attachment to abiotic surfaces, (iii) High abundance of proteases and proteolytic molecules, hindering cell-cell adhesion, and cellular aggregate formation in the EPS matrix, and (iv) Ecotin mediated inhibition of proteases that are necessary during biofilm dispersal. We reasoned that the lower abundance of redox homeostasis proteins and ferritin-like molecules in the Δ*yjiY* supernatant might facilitate the accumulation of toxic oxidative species, causing cellular stress and toxicity. In this light, these oxidative stress relieving proteins can be exploited as potential druggable targets to inhibit *Salmonella* biofilm initiation.

## Materials and Methods

### Bacterial strains

All *Salmonella* Typhimurium strains used in this study are listed in the following section with their genetic descriptions. *Salmonella enterica* serovar Typhimurium strain 14028S was used as the wild type strain, and was also the parental background for all the mutant strains used in this study, i.e. Δ*yjiY, ΔcsgD, ΔfliC* and Δ*fliC ΔfljB*. All strains were grown and maintained in Lennox broth (LB; 0.5% NaCl, 1% casein enzyme hydrolysate and 0.5% yeast extract) at 37°C under shaking conditions. STM GFP, STM mCherry (RFP-STM-WT) and STM Δ*yjiY:yjìY* were cultured in Lennox broth with 50 μg/ml Ampicillin at 37°C in shaking condition. *E.coli* DH5α, *Staphylococcus aureus, Pseudomonas aeruginosa* and *Klebsiella pneumoniae* were grown in Lennox Broth at 37°C in shaking condition.

List of strains used in this study.

| Strain name | Description | Reference |
| --- | --- | --- |
| *S. Typhimurium*<br>ATCC 14028S<br>(STM_WT) | Wild type (WT) | Kind gift from Prof. M. Hensel (Division of Microbiology, University of Osnabrück, Germany) |
| $\Delta yjiY$ | Isogenic knockout strain for the gene *yjiY*; Kan<sup>r</sup> | Garai *et al.* (2016) |
| $\Delta csgD$ | Isogenic knockout strain for the gene *csgD*; Chl<sup>r</sup> | Srinandan *et al.* (2015) |
| $\Delta fliC$ | Isogenic knockout strain for the gene *fliC*; Chl<sup>r</sup> | Garai *et al.* (2016) |
| $\Delta yjiY$-pQE60-*yjiY* | Isogenic complement strain for $\Delta yjiY$ expressing *yjiY* under the promoter of T5 present in the plasmid pQE60; Amp<sup>r</sup> | Garai *et al.* (2016) |
| $\Delta fliC \Delta fljB$ | Isogenic double knockout strain for *fliC* and *fljB*; Kan<sup>r</sup> Chl<sup>r</sup> | Karmakar *et al.* (2019) |
| STM mCherry<br>(RFP-STM-WT) | STM 14028s wildtype strain carrying<br>pFPV-mCherry plasmid, Amp<sup>r</sup> | This study |
| STM GFP | *S. Typhimurium* rpsM promoter<br>driving gfpmut3 expression through<br>pFPV25.1 plasmid, Amp<sup>r</sup> | This study |

### Growth conditions for biofilm formation

LB without NaCl, i.e. 1% casein enzyme hydrolysate and 0.5% yeast extract, was used as biofilm media. Overnight cultures grown in LB were subcultured in 2 ml biofilm medium at the dilution of 1:100 in a flat-bottom 24-well polystyrene plate (Tarsons) and incubated at 28°C for 72 hours without shaking. All images of biofilm in the form of a pellicle were taken with a digital camera (Olympus Stylus VH-520).

### Preparation of cell free supernatant

To 2 mL of biofilm media, 1:100 dilution of overnight grown culture was added in flat-bottom 24-well polystyrene plate (Tarsons). The plate was incubated at 28°C without shaking for 72 hours. After 72 hours, the media from each well was collected, centrifuged twice at 10000 rpm for 15 minutes, the supernatant was collected and filtered using a 0.2 μm filter and stored at −20°C. To collect supernatant from M9 (0.5% glucose) media, the cultures were grown at 37°C in shaking conditions. After 72 hours, the supernatant was collected as described above.

### Bacterial biofilm inoculation with culture supernatant

To test the effect of the 3 day culture supernatant (chemically treated and untreated) on biofilm formation, 20 μl of STM WT overnight culture and indicated volume (1% v/v and/or as indicated in the figures) of the supernatants (WT supernatant, Δ*yjiY* supernatant, Δ*yjiY:yjiY* supernatant and Δ*csgD* supernatant) was added to 2 ml of biofilm media, and incubated under biofilm inducing condition. **Cell pellets** were collected from 3 day old biofilm culture by harvesting the contents of the flat-bottom 24-well polystyrene plates (Tarsons), centrifuging at 10000 rpm for 15 minutes. The cells were resuspended in fresh biofilm inducing media and added at the concentration of 0.5×10^7^ CFU/ml of biofilm media and 20 μl of STM WT overnight culture, and incubated under biofilm inducing condition. For **co-culture** experiments, the overnight cultures of the strains were added in 1:1 ratio to 2 ml of biofilm media and incubated under biofilm inducing condition. **Cell lysates** were prepared by adding 30 μl 1x TME buffer (25 mM Tris pH 8.0, 1 mM EDTA pH 8.0, 2 mM ß-mercaptoethanol) and sonicating the bacterial samples (STM WT, STM Δ*yjiY*, and STM Δ*yjiY:yjiY)*. The samples were centrifuged at 12000 rpm for 10 minutes and 2 μl supernatant was used for inoculation along with 20 μl STM WT overnight culture and 2 ml of biofilm media and incubated under biofilm inducing condition.

### Crystal Violet staining

To quantify the biofilm at the solid-liquid interface, crystal violet (CV) staining was carried out as mentioned in Chandra *et al*(14). Briefly, the protocol followed for biofilm formation was the same as mentioned above. After 72 hours, the media in each well was discarded and the plates were thoroughly washed with RO water to remove all planktonic cells. The plates were then dried and 2 ml of 1% CV was added into each well. After 15 minutes, the CV was removed, and the excessive stain was thoroughly rinsed with RO water. The stained biofilm was destained with 70% ethanol and the intensity of color of the destained solution was quantified at OD595 in Tecan plate reader (Infinite Pro 200). The absorbance was plotted in GraphPad Prism 6 and significance values determined using Student’s t-test or two-way ANOVA.

### Confocal microscopy

Sterile square coverslips (18 mm) were placed in flat-bottom 12-well polystyrene plate (Tarsons). Cultures were inoculated for biofilm formation as mentioned previously. After 72 hours, biofilm appeared on the coverslip at the liquid–air interface, in the form of a thin line spanning the width of the coverslip (18 mm). The coverslip was washed thoroughly with water to remove planktonic cells and stained with Congo red (20 mg/ml in water) for 20 min at room temperature. After washing with water, the coverslip was mounted on a slide and imaged for biofilm distribution, with a laser scanning confocal microscope (Zeiss LSM 710) using a 40x objective. Z stacks were taken to generate a three-dimensional image. The MFI of the images were calculated using the ImageJ software. The MFI and thickness of the biofilm were plotted using GraphPad Prism 6. Single layer of cells were imaged and cell length of ~1000 cells from each coverslips was measured using ImageJ.

### Glass bead assay

To test the strength of the pellicle at the air-liquid interface, medium sized (0.5mm to 1mm) glass beads were added one by one onto the pellicle. The initial weight of the glass beads was noted, and the number of glass beads added until the pellicle just collapsed was noted down and plotted using GraphPad Prism 6.

### Scanning electron microscopy

Biofilm was allowed to form on coverslips as mentioned in the previous section. After thorough washing with water, the sample was fixed in 2.5% gluteraldehyde for 48 hours at room temperature. Excess gluteraldehyde was removed by washing with water and the sample was dehydrated by gradient washes in increasing concentrations of 30%, 50%, 75%, 85% and 95% ethanol. The coverslips were then air dried under vacuum before coating with gold (JOEL-JFC-1100E ion sputtering device) for imaging by scanning electron microscope. For checking flagellar morphology, 20 μl of STM WT overnight culture and 20 μl of supernatant treated STM WT were smeared on autoclaved 18mm coverslips, air dried, and processed using the abovementioned protocol. Flagellar structure was imaged using field emission-SEM (FEI Sirion, Eindhoven, The Netherlands) scanning electron microscope.

### Atomic force microscopy

Sterile 18 mm square coverslips were placed in flat-bottom 12-well polystyrene plates (Tarsons) and biofilm inoculation was done. After 72 hours, the coverslips were removed and washed with sterile MilliQ water. The coverslips were dried and AFM analysis was done using XEISS AFM systems and was analyzed using XEI software.

### Supernatant conditioning

The supernatant was treated with different concentrations of EDTA (2.5 mM, 5 mM, 7.5 mM and 10 mM) and incubated at 65°C for 1 hour. To 200 μl of supernatant, 10 μl of proteinase-K (NEB, stock 20 mg/ml) was added and incubated at 37°C for 1 hour. The proteinase was inactivated with 0.5 mM PMSF and incubated at 28°C for 1 hour. The supernatant was heated to 37°C (60 min), 65°C (15 min) or 95°C (15 min) for indicated time and immediately frozen at −20°C. The pH of the biofilm inducing media as well as that of the supernatant were adjusted using concentrated HCl and 10N NaOH to obtain pH of 4, 7.4, and 9. The supernatants were also treated with 10 μl RNase (stock 1 mg/ml) for 1 hour at 37°C. Post treatment, the supernatants were used to inoculate biofilm in order to check the activity of the inhibitory molecule(s).

### Preformed biofilm disruption

STM WT was allowed to form biofilm as described earlier. 72 hours post inoculation, the formed biofilms were treated with 20, 50 or 100 μl (1% (v/v), 2.5% (v/v) or 5% (v/v), respectively) supernatants and was incubated at 28°C for 72 hours and biofilm was quantified using CV staining.

### Recovery of biofilm formation

STM WT was allowed to form biofilm in the presence of supernatants as described earlier. 72 hours post inoculation, the cells were harvested by centrifuging the culture at 10000 rpm for 15min. 20 μl of these treated or untreated cells were inoculated in fresh biofilm media (with and without supernatants for retreatment or recovery, respectively) and incubated at 28°C for 72 hours and biofilm was quantified using CV staining.

### Supernatant concentrating and separation based on molecular weight

The supernatants from biofilm culture were harvested and filter sterilized as previously mentioned. 4 ml of the supernatants were transferred to the Amicon ultra filter device (Amicon® Ultra-4 centrifugal filter device, 3k MWCO, UFC800324) and centrifuged at 4000g for 30 min in a swing bucket rotor. The concentrated solute from the bottom of the filter device was collected by inserting a pipette.

### Quantitative RT PCR

STM WT was allowed to form biofilm in the presence or absence of supernatants as described earlier. After 72 hours, the biofilm population as well as the planktonic cells were harvested by thorough pipetting and centrifugation. From these cells, RNA was isolated by the TRIzol method (Takara). cDNA was synthesized with reverse transcriptase (GCC Biotech). Quantitative PCR was carried out using SYBR Green Q-PCR kit (Takara).

List of oligonucleotides used in this study.

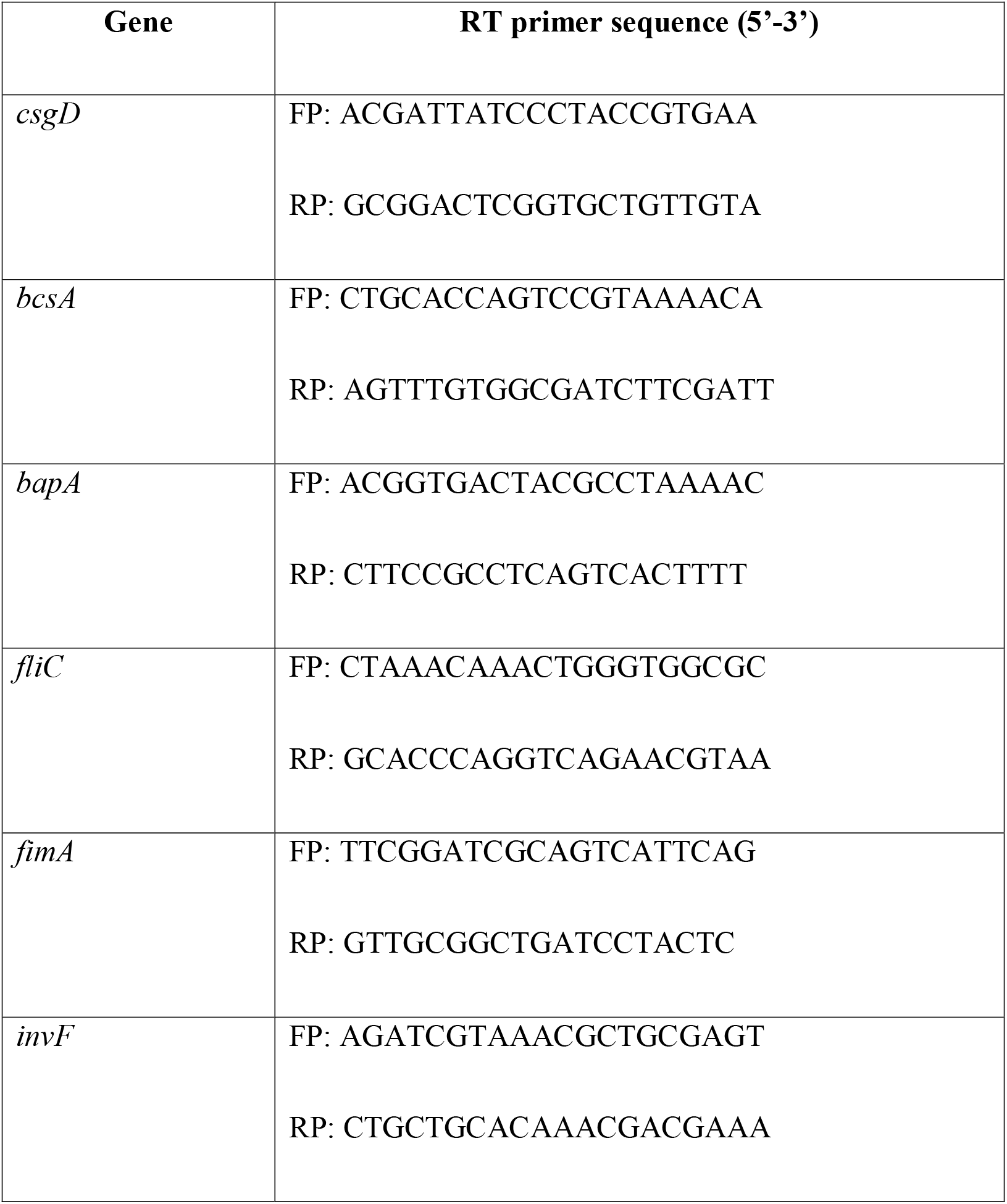
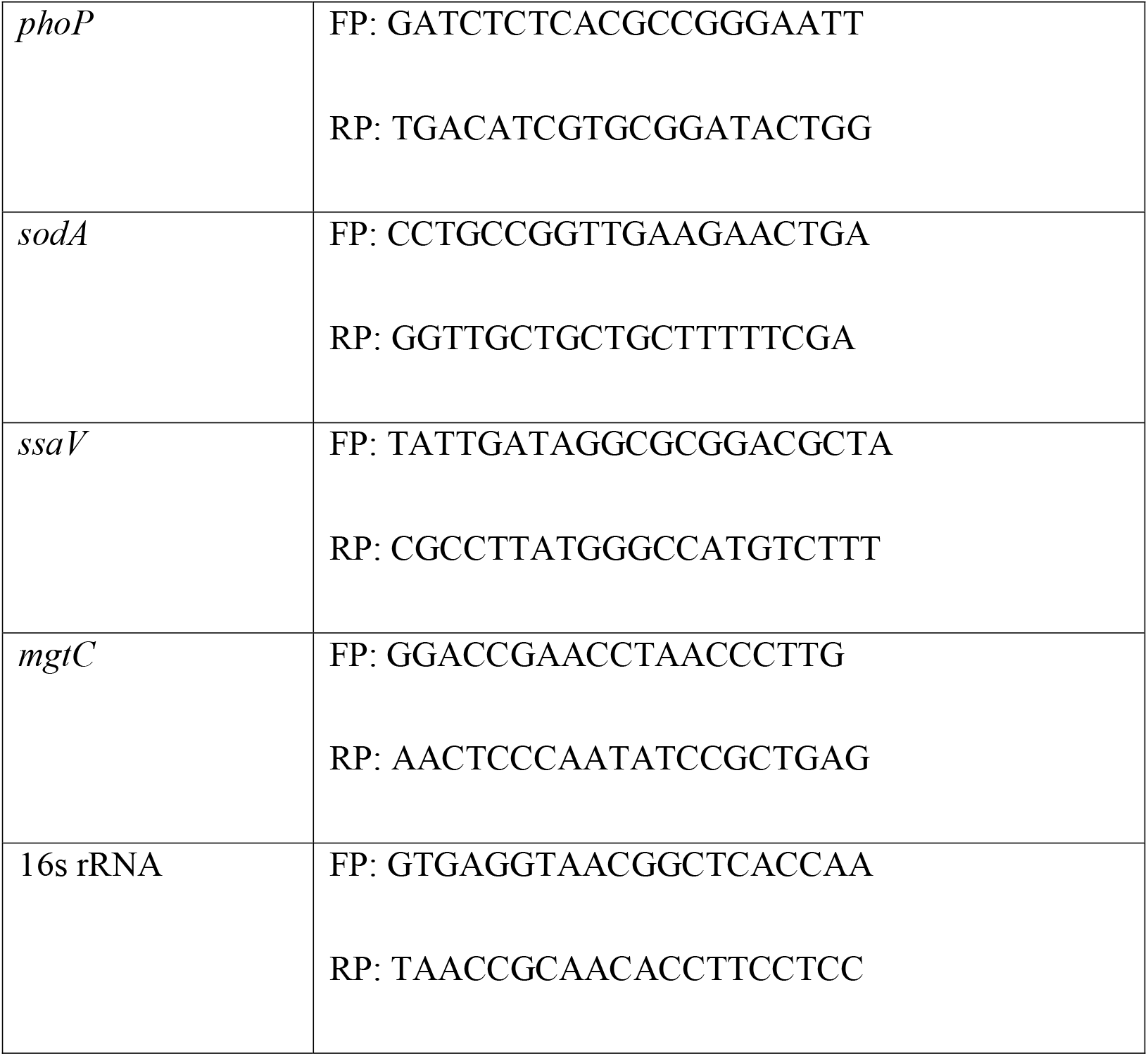

### *In vitro* motility assay

2 μl of bacterial samples (treated or untreated) were spotted onto the 0.3% agar plates supplemented with 0.5% yeast extract, 1% casein enzyme hydrolysate, 0.5% NaCl and 0.5% glucose (swim agar plates) or 0.5% agar plates supplemented with 0.5% yeast extract, 1% casein enzyme hydrolysate, 0.5% NaCl and 0.5% glucose (swarm agar plates). The plates were incubated at 37°C and images were taken every 2 hours using a digital camera (Olympus). The diameters of the motility halos were measured using ImageJ. At least five replicate plates were used for each condition, and statistical significance was calculated using Student’s *t*-test.

### Transmission electron microscope

Flagella and fimbriae were visualised using the protocol described in Garai et al 2016(9). Briefly, overnight STM WT were inoculated with or without supernatants in biofilm media and kept under biofilm inducing condition for 72 hours. After 72 hours, both biofilm and planktonic cells were collected and fixed with 2.5% glutaraldehyde overnight at 4°C. Similarly fixed overnight cultures of STM WT and STM Δ*fliC ΔfljB* were used as positive and negative control, respectively. 2 μl of the cell suspension was added on copper grid, air dried, stained with 1% uranyl acetate for 30 sec, and visualised under transmission electron microscope.

### *In vitro* intracellular survival assay

Int407 cells (ATCC® CCL-6™) were seeded at a density of 2×10^5^ cells per well in flat-bottom 24-well polystyrene plates (Tarsons). 72 hours old supernatant treated or untreated biofilm cultures were sub-cultured, grown to mid-log phase and used for infection of Int407 at MOI 10. The plates were incubated at 37°C with 5% CO_2_ for 20 min to allow infection of the cells. The cells were incubated for an hour in 100 μg/ml Gentamicin. The cells were then washed, and grown in DMEM containing 25 μg/ml Gentamicin. After 1h, 2h and 16h, the cells were lysed with 0.1% Triton-X100 and bacterial load was estimated by plating on SS-agar plates.

### *In vivo* virulence in *C. elegans* N2 worms

STM-RFP bacteria were treated with the supernatants, and kept for biofilm formation for 72 hours. After that, STM-RFP cells were harvested as mentioned before, and 10^7^ CFU were seeded on NGM plates, incubated for 12 hours at 37 °C. *E. coli* OP50 fed *Caenorhabditis elegans* N2 young adults were harvested in M9 buffer, counted and approximately 150 worms were fed with supernatant treated or untreated 10^7^CFU of STM WT RFP bacteria for 12 hours. The worms were then harvested in 1ml M9 buffer, washed 6-8 times to remove extracellular *Salmonella*. 10 worms from each sample were transferred to sterile M9 buffer containing 0.5mm sterile glass beads. The worms were lysed, and the gut content of the worms was plated on SS-agar plates. ~20 worms were taken for laser scanning confocal microscopy analysis using Zeiss LSM 880 with Multiphoton mode. ~50 worms were transferred to *E. coli* OP50 plate and fed for another 12 hours. After 12 hours, the worms were sampled in a similar manner for CFU analysis and imaging.

### SDS-PAGE and colloidal Coomassie Blue staining

The proteins in the supernatants were quantified by Bradford’s assay and equal quantity of the STM WT and STM Δ*yjiY* supernatants were loaded onto the SDS-PAGE gel (5% stacking, 10% resolving). The gel was then stained with colloidal Coomassie Brilliant Blue R-250 for 3 days and then washed with water to remove excessive stain and visualize the band.

### LC QTOF MS/MS analysis of the supernatants

Total protein from the 10-fold concentrated supernatants was subjected to in-solution trypsin digestion for LC Q-TOF MS analysis. Briefly, 200 mg of total protein was resuspended in 200 μl of 50 mM Ammonium bicarbonate. To this, 10 mM DTT was added, and incubated at 56°C for 30 min, followed by alkylation with 20 mM iodoacetamide and incubation at 37°C for 30 min in dark. For protein digestion, trypsin solution was added to the sample to a final protease to protein ratio of 1:50 (w/w) and incubated at 37°C for 16 hours, with frequent shaking. The reaction was stopped by adding 10 μl of 0.1% formic acid, and samples were stored at −20°C until loaded on the Agilent AdvanceBio Peptide Map column (2.1 ×150 mm, 2.7μ). 20 μl of the digested samples were injected to the column, and analysed at a flow rate of 0.4 ml/min using 0.1% formic acid in water as mobile phase A and 0.1% formic acid in acetonitrile as mobile phase B.

All the samples were analyzed using Agilent 1290 Infinity II LC System coupled with Agilent AdvanceBio Q-TOF (6545XT). The MS and MS/MS scan were acquired in the positive mode and stored in centroid mode. The MS data acquisition parameters were as follows-Vcap was set at 3500V, drying gas flow rate and temperature was set at 13 ml/min and 325 °C, respectively. A collision energy with a slope of 3.6 V/100 Da and an offset of 4.8 V was used for fragmentation. The Precursor ion data were captured in a mass range of *m/z* 200–2700 and fragment ions data were acquired between *m/z* 50–2800. The raw data were analyzed using MaxQuant software (v1.6.2.10) and processed through MS excel sheet. The database analysis was performed against *Salmonella* Typhimurium proteome in UniProt (Proteome ID: UP000002695). Following search parameters were used for the database analysis: Precursor mass tolerance: 10ppm, fragment mass tolerance: 40ppm, fixed modifications: carbamidomethyl (C), variable modification: oxidation.

### Cloning, expression and purification of ecotin, HNS and NusG

*S*. Typhimurium 14028S Ecotin (STM14_2792), HNS (STM14_2116) and NusG (STM14_4985) were cloned with 6x His tag in pET15b vector using Gibson assembly. Positive clones were confirmed by PCR and transformed in *E. coli* BL21 (DE3) pLysS strain. *E. coli* BL21 (DE3) cells bearing expression plasmids were grown overnight at 37°C in LB media containing 50 μg/ml chloramphenicol and 100 μg/ml ampicillin. 1% of the primary inoculum was added to 500ml of LB media containing 100 μg/ml ampicillin and grown at 37°C till the culture attained an OD_600_ of ~ 0.8, and induced with 1mM isopropyl β-D-1-thiogalactopyranoside (IPTG) at 30°C for 6 hours. The cells were harvested by centrifugation, and the pellet was resuspended in 20ml ice-cold resuspension buffer (0.2M Tris (pH 7.5), 0.5mM EDTA, and 1mM PMSF). Further, the cells were lysed by sonication on ice, followed by centrifugation at 14000g. The protein was purified from the soluble fraction of the lysate by affinity chromatography using Ni-NTA column. Protein was eluted with linear gradient of 0.1-1M imidazole in 50mM Tris (pH 8.0). Elute fractions containing protein of interest were pooled, concentrated and subjected to purification by size exclusion chromatography (SEC) on Superdex 75 Increase 10/300GL column (with a flow rate of 0.5 ml/min and 500 μl sample injection volume) equilibrated with 50mM Tris (pH 8.0). The peaks obtained from the chromatogram were analyzed by subjecting the collected fractions (which were pooled and concentrated after SEC) to SDS-PAGE, to obtain the protein of interest in its desired monomeric form. The protein purity was also confirmed by MALDI-TOF.

List of oligonucleotides used for cloning.

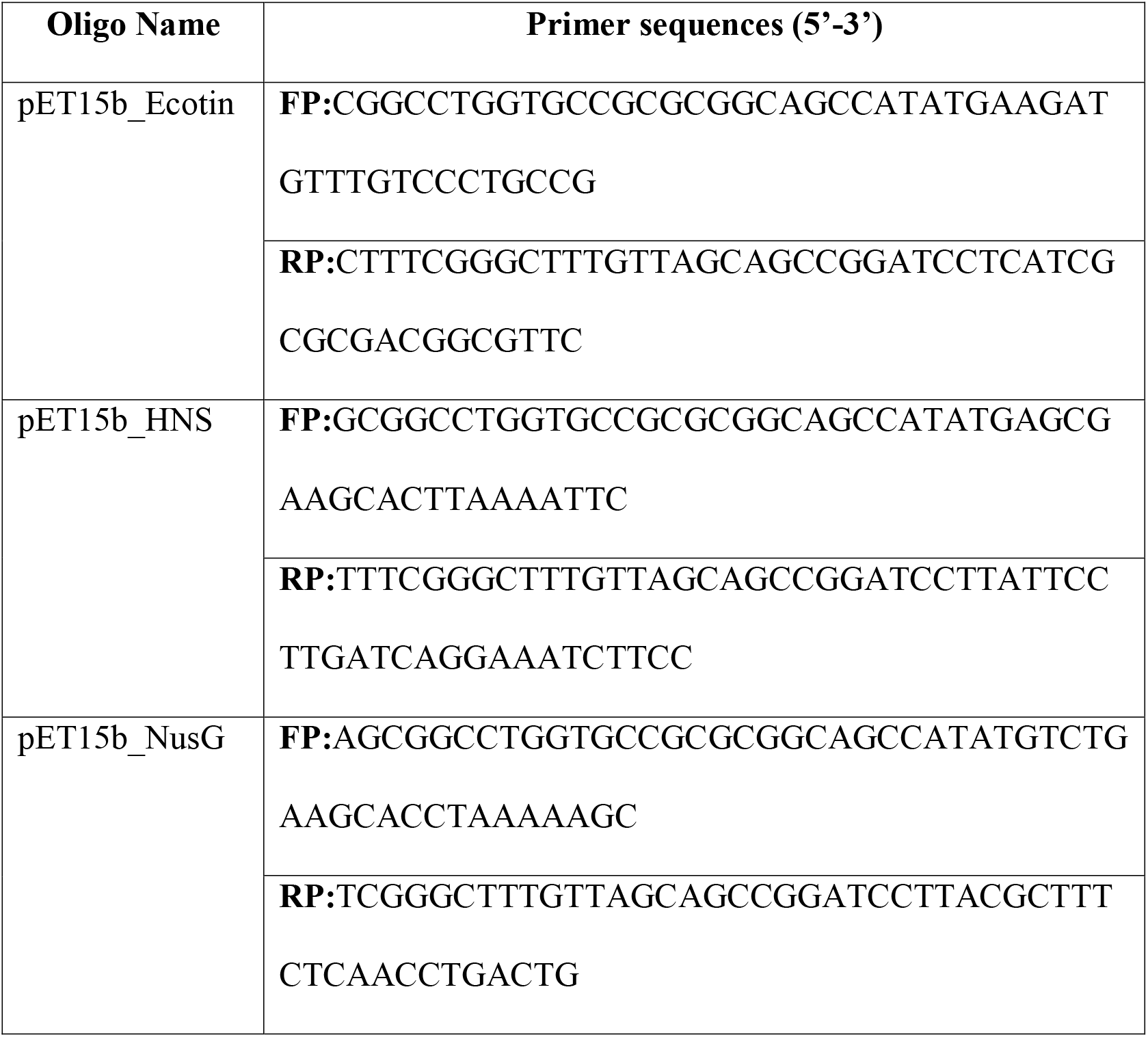

### Statistical analysis

Statistical analyses were performed with GraphPad Prism software. Student’s t test and one-way or two-way ANOVA tests were performed as indicated. The results are expressed as mean + SD or mean + SEM. Group sizes, experiment number, and p values for each experiment are described in the figure legends.

## Acknowledgements

We thank the electron microscopy facility of IISc, AFMM facility of IISc, Atomic force microscopy facility of BSSE, IISc and confocal microscopy facility and real-time facility of Dept. of Microbiology and Cell Biology, IISc. We thank Mr. Priyabrata (TH lab, MRDG, IISc) for the helping with the TEM sample preparation. We thank Dr Shahbaz and Dr Jayantika (RV lab, MBU, IISc) for helping with cloning, protein expression and purification. We are very grateful to Dr Kapudeep and Ms. Meghanashree for their valuable inputs.

## Author Contribution

KC and DC conceived the study and designed experiments. KC and PM performed experiments. SK performed and UST provided valuable inputs for the LC QTOF MS/MS experiment. KC analyzed the data, prepared the figures and wrote the manuscript. DC supervised the work. All the authors read and approved the manuscript.

---

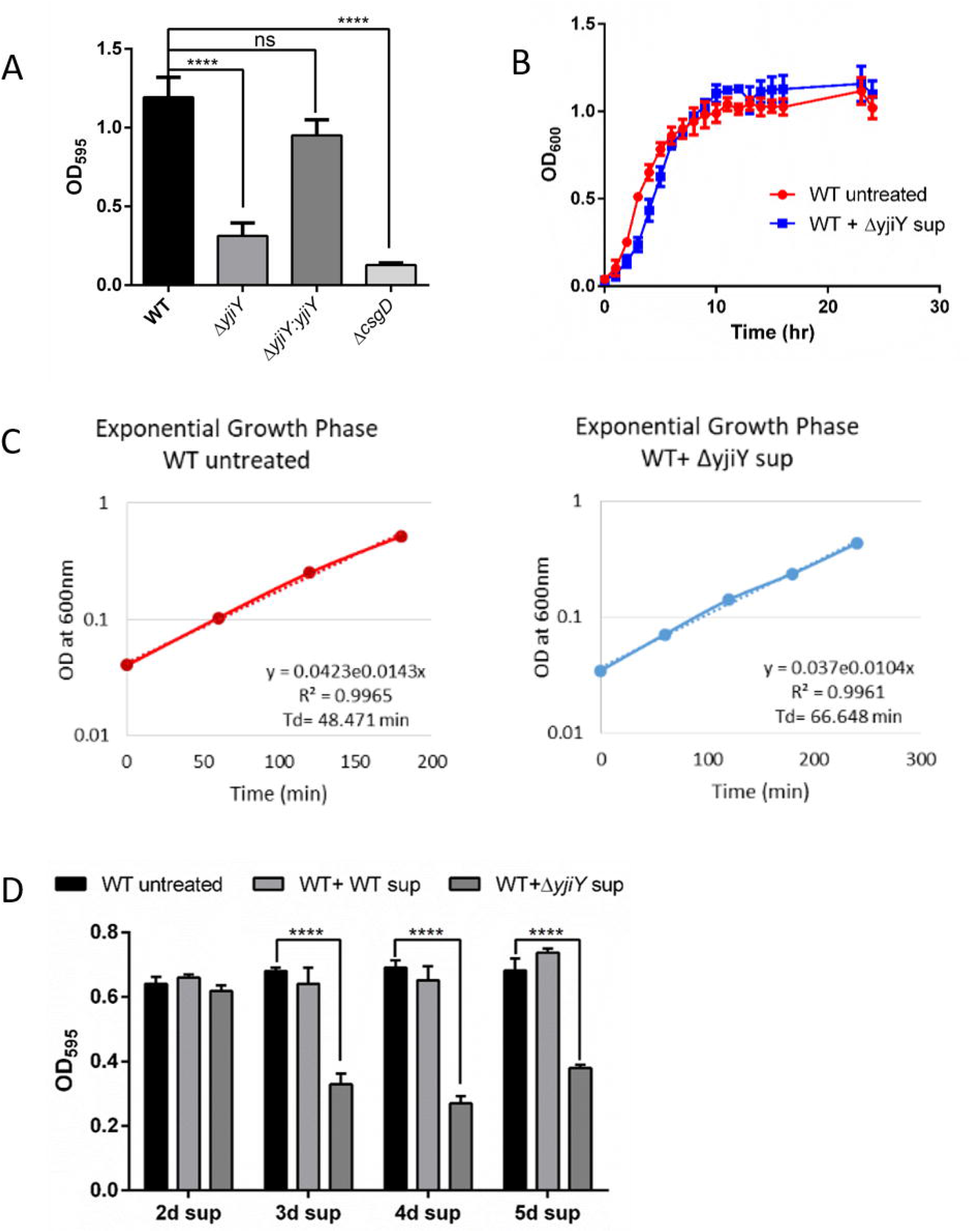

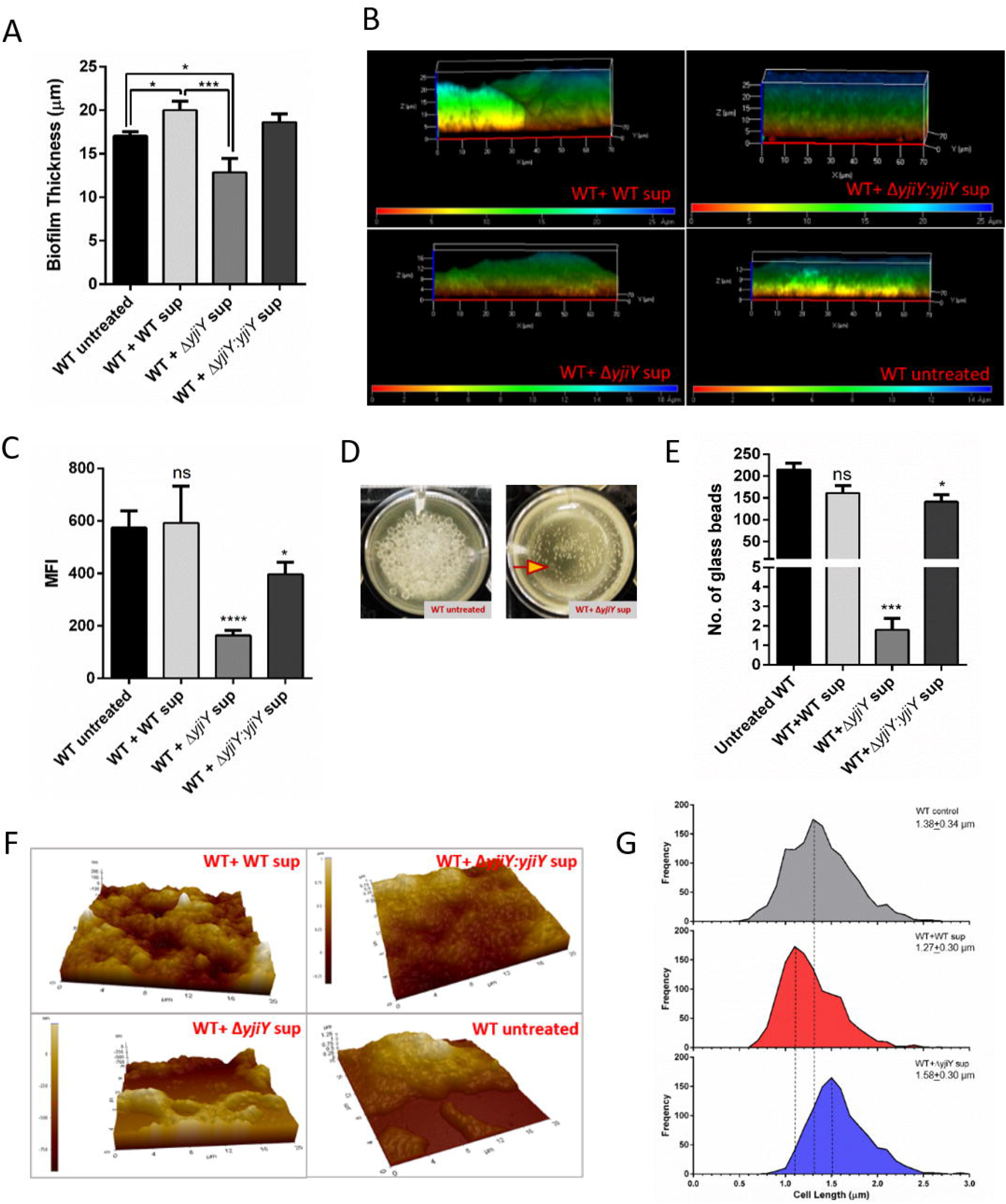

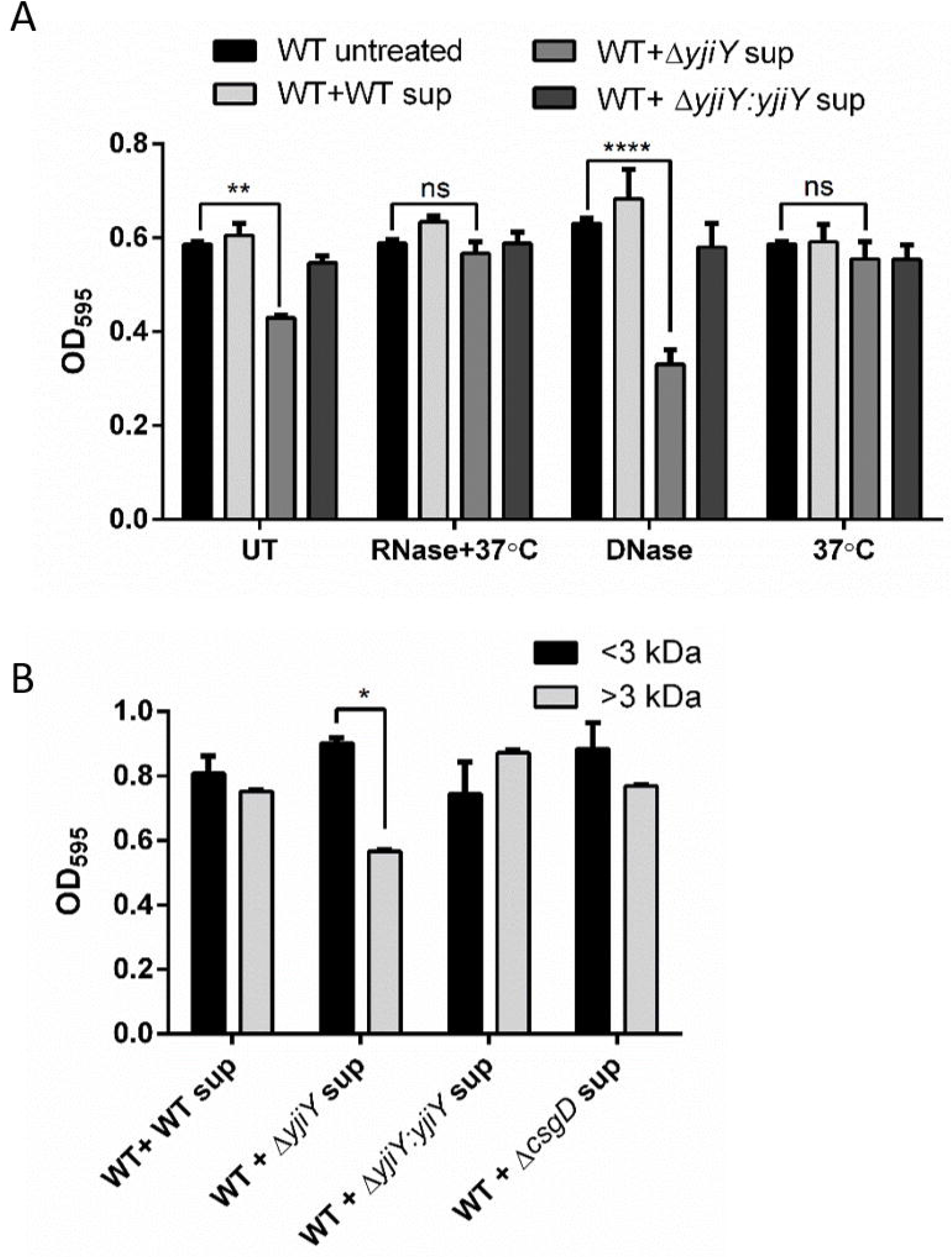

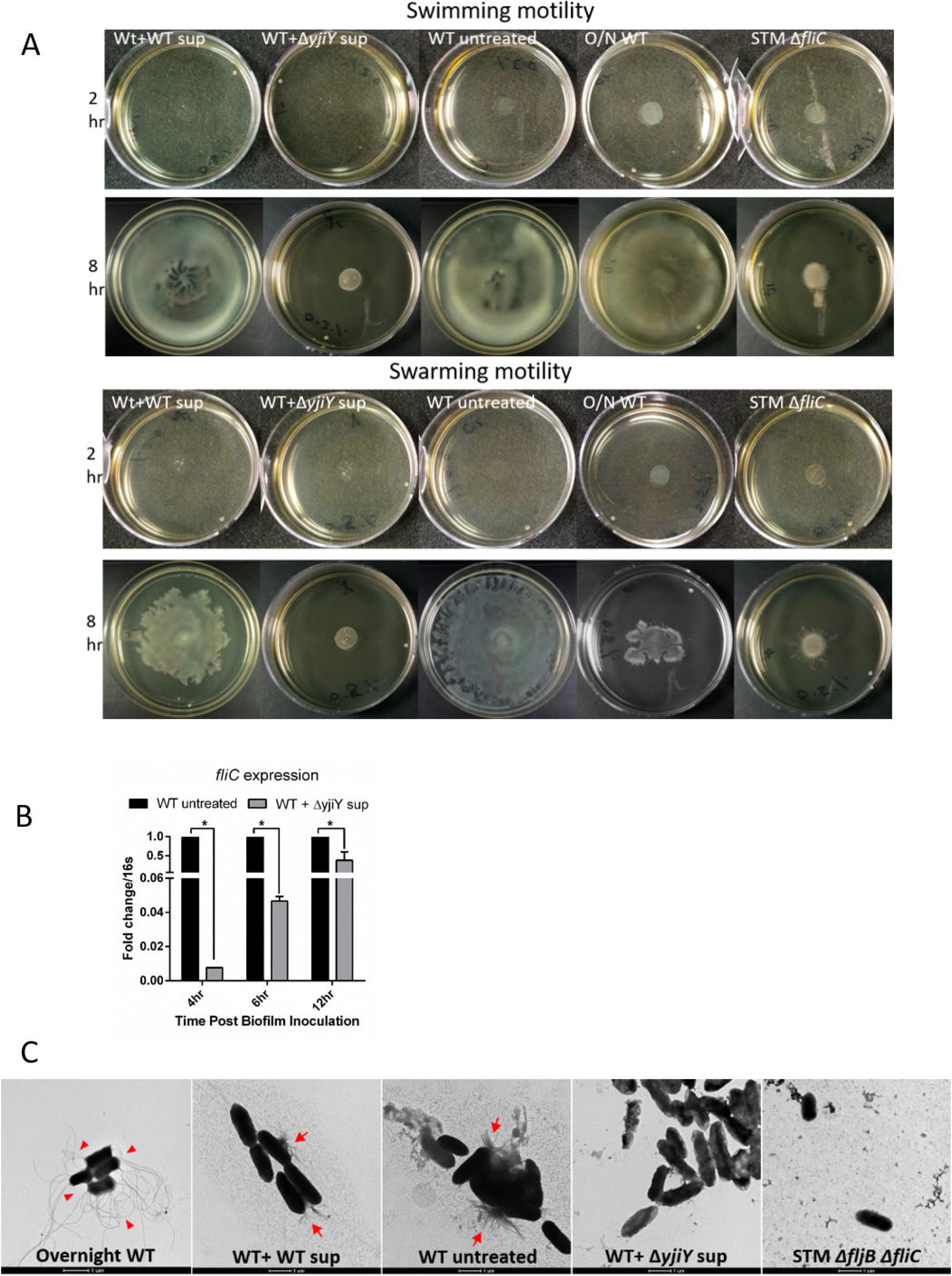

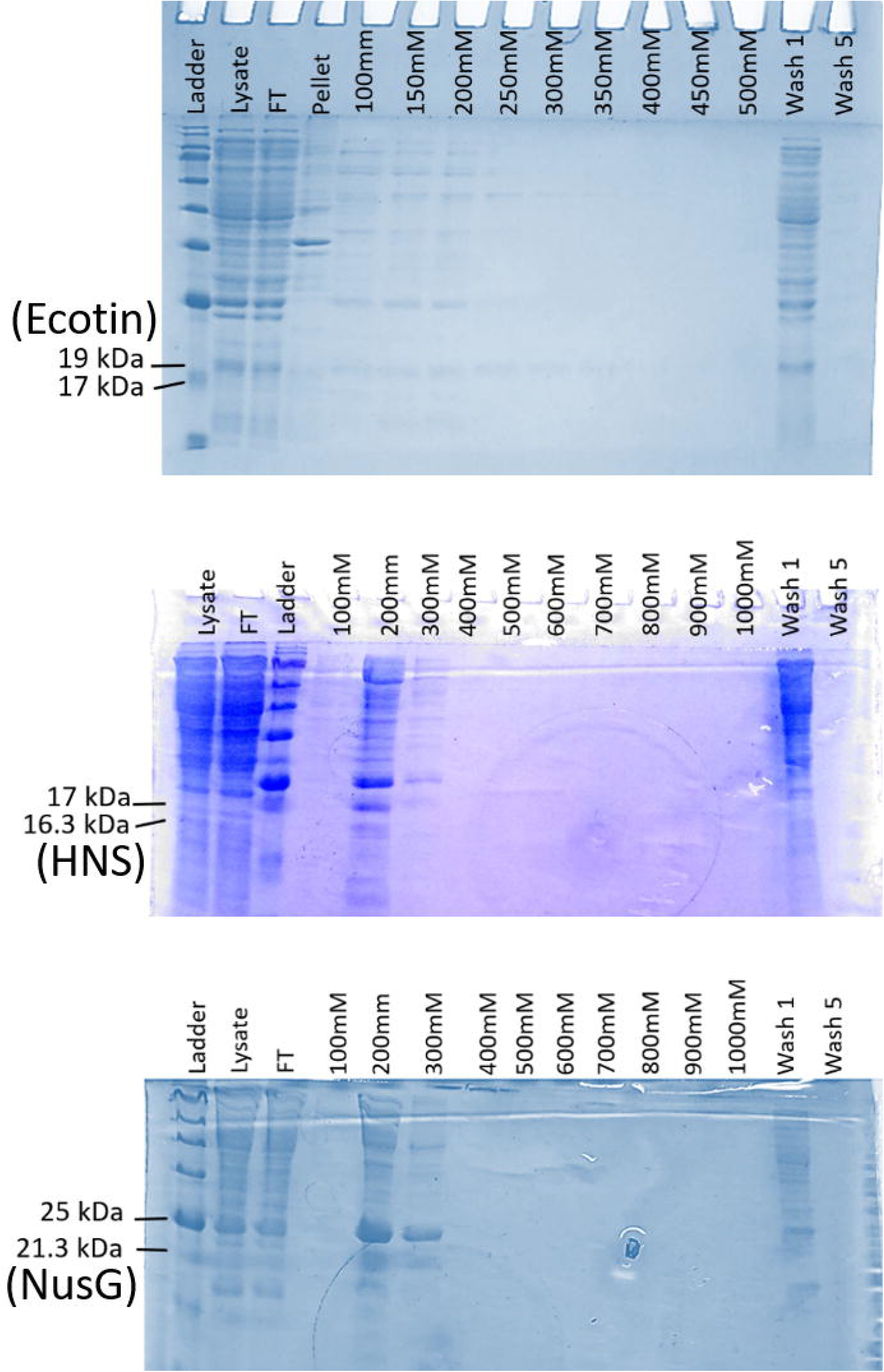

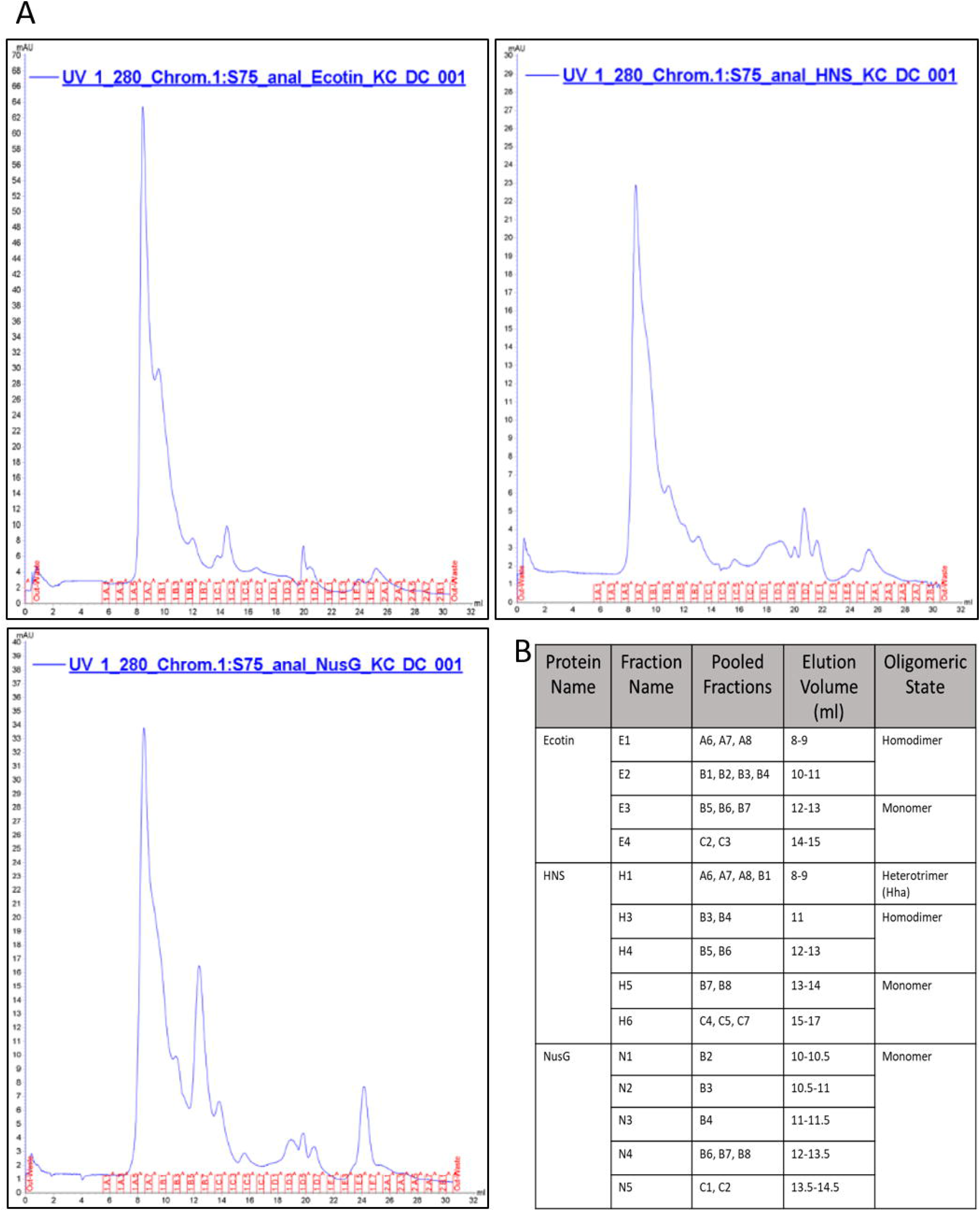

