## Supplementary Figure legends and tables for "Global transcriptional regulation by cell-free supernatant of *Salmonella* Typhimurium peptide transporter mutant leads to inhibition of intra-species biofilm initiation"

Dipshikha Chakravortty

**Keywords**: *Salmonella* Typhimurium, Biofilm, flagella, H-NS, EPS, yjiY, oxidative stress

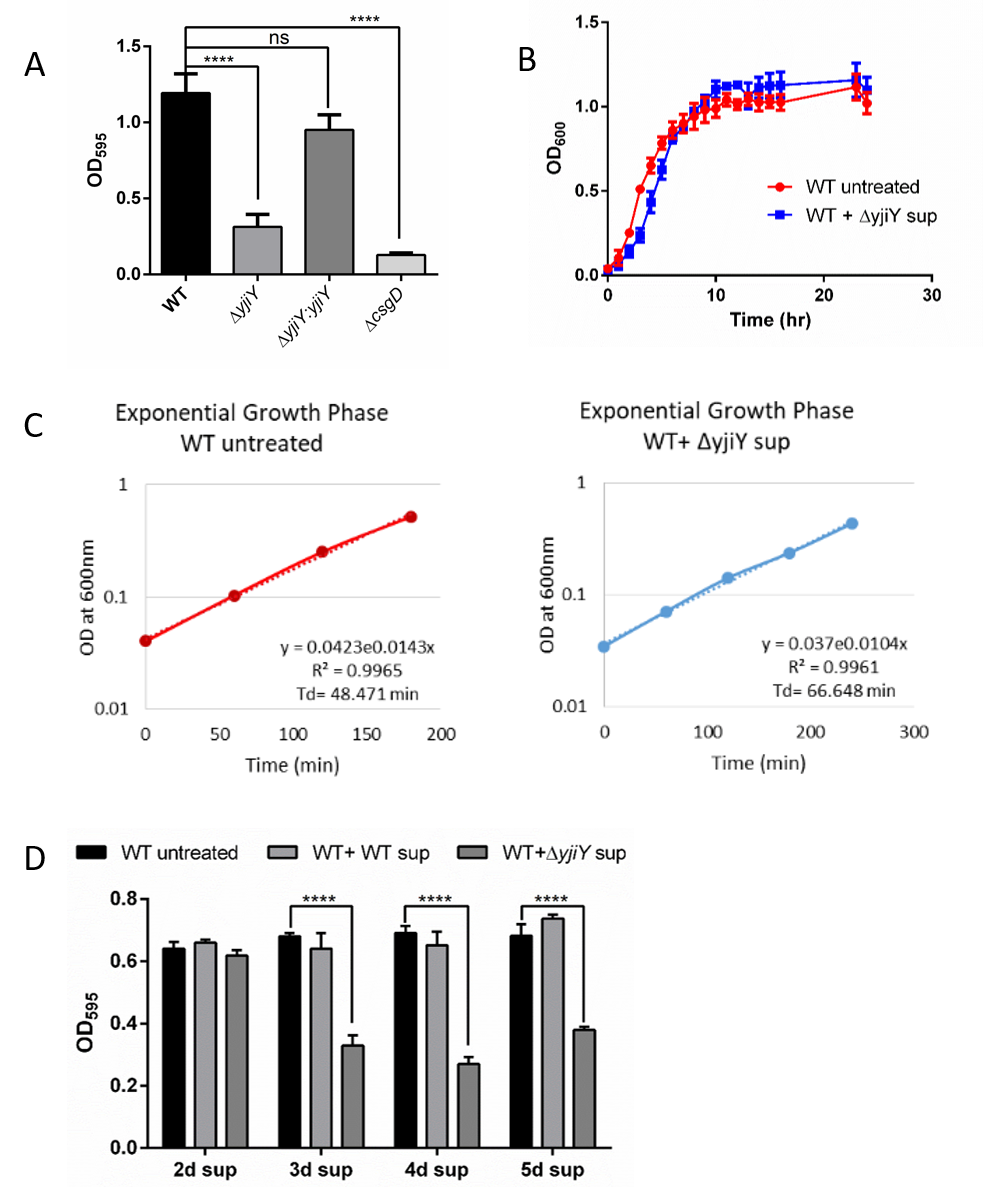

**Fig S1. *Salmonella* *ΔyjiY* supernatant lacks any bactericidal activity, and significant biofilm inhibitory activity is present in 3day old supernatant**

A. Biofilm formation ability of the different strains used in this study was checked (Data are presented as mean ± ­SEM of 5 independent experiments). B. The growth of STM WT bacteria was checked in presence and absence of *ΔyjiY* supernatant (Data are presented as mean ± ­SEM of 3 independent experiments). C. Graphs showing exponential phase of growth of untreated WT (left panel: red line) and *ΔyjiY* supernatant treated WT (right panel: blue line) cells. (Data are presented as mean of 3 independent experiments). D. Biofilm inhibition activity of *ΔyjiY* supernatant, that was collected on different days of biofilm inoculation with pure culture, was checked (Data are presented as mean ± ­SEM of 5 independent experiments).

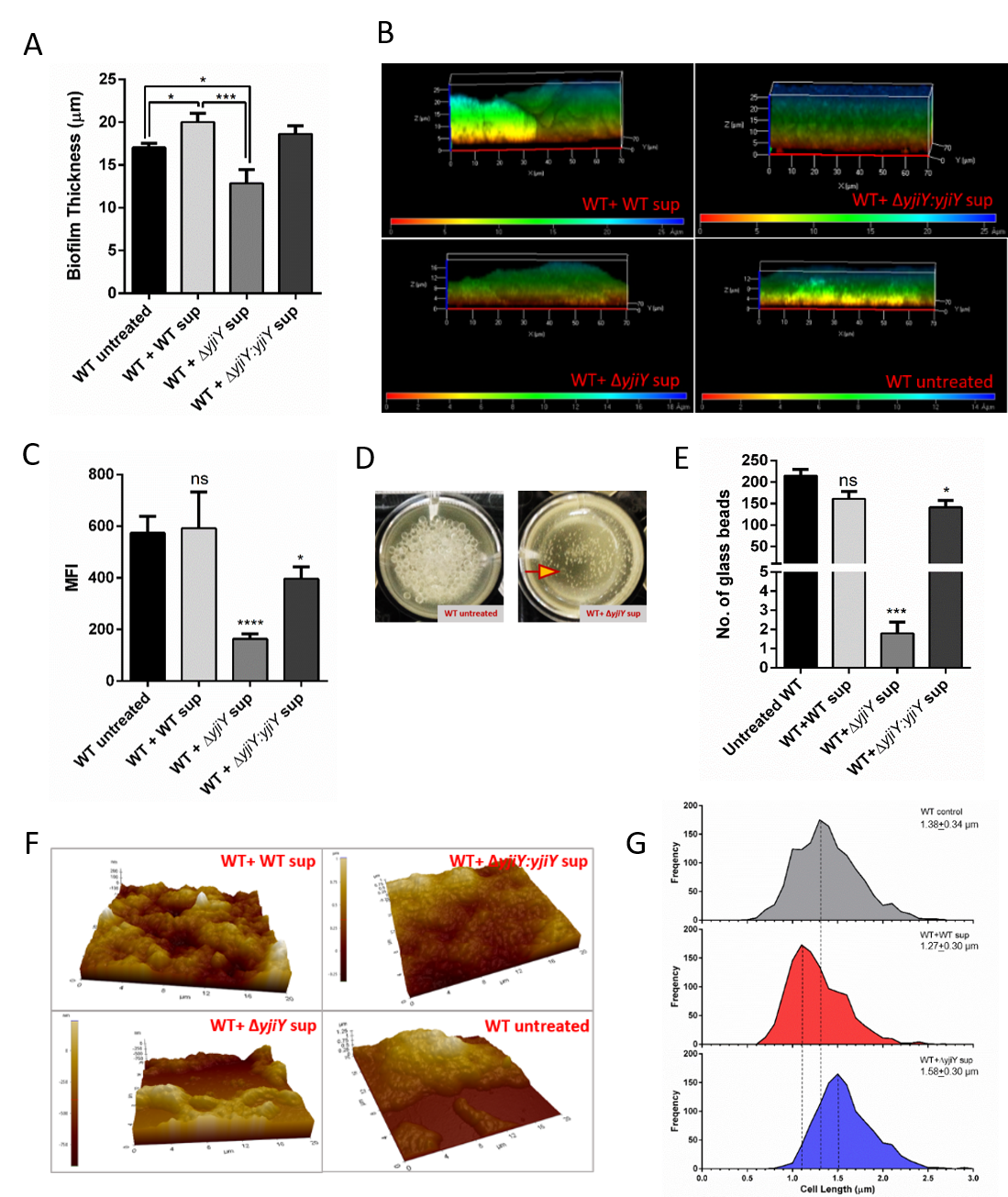

**Fig S2. Δ*yjiY* supernatant makes the biofilm thin, fragile as well as it modifies cell length**

A. The thickness of the biofilm formed on glass coverslips was measured using Zen (black edition) and plotted using GraphPad Prism 6 (Data are presented as mean ± ­SEM of 5 independent experiments). B. Representative CSLM images of the Congo red stained biofilm formed on glass coverslip, depth coding showing a reduced thickness of biofilm after Δ*yjiY* sup treatment (Representative image from 5 independent experiments). C. Median fluorescence intensity (MFI) of the Congo red stained biofilm was measured using ZEN (Black) software and MFI values were plotted with GraphPad Prism 6 (Data are presented as mean ± ­SEM of 5 independent experiments). D. Representative image of the strong and fragile biofilm formed without or with Δ*yjiY* sup. Yellow arrow shows presence of only one glass bead at the bottom of the well (Representative image from 3 independent experiments). E. Number of glass beads required to sink the biofilm pellicle to the bottom was counted and plotted using GraphPad Prism (Data are presented as mean ± ­SEM of 3 independent experiments). F. Representative Atomic Force Micrograph images of biofilm on the glass coverslip showing the absence of characteristic dome shaped structure of biofilm with Δ*yjiY* supernatant treatment (Representative image from 2 independent experiments). G. Frequency distribution of the cell length with different supernatant treatment showing a shift towards longer cell length with Δ*yjiY* sup treatment (Data are presented as mean ± ­SEM of 3 independent experiments, length of approximately 1200-1400 cells from each treatment were measured). One-way ANOVA was used to analyze the data, p values ****<0.0001, ***<0.001, **<0.01, *<0.05.

**Fig S3. The active component(s) is/are not RNA or DNA and the components are larger than 3kDa in size**

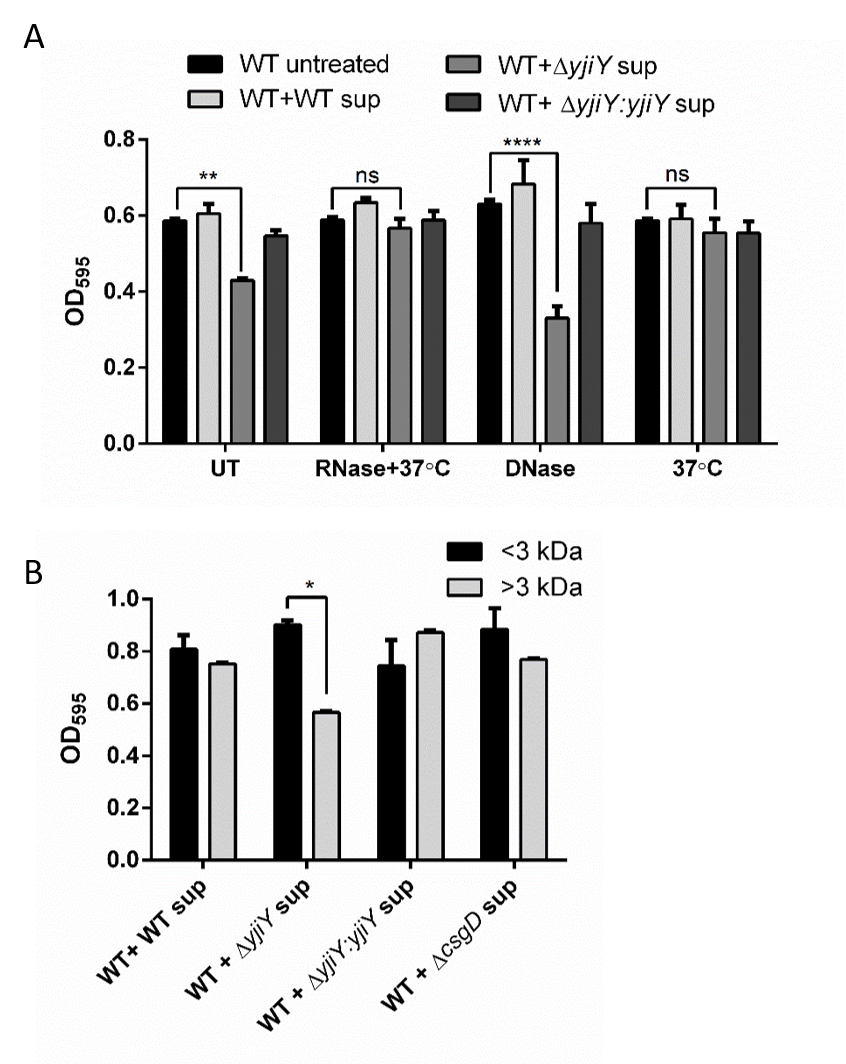

A. The supernatants were treated with RNase at 37˚C for 1 hour, as well as with DNase for 1 hour at 65˚C (Data are presented as mean ± ­SEM of 3 independent experiments). One-way ANOVA was used to analyze the data, p values ****<0.0001, **<0.01. (UT- Untreated sup treated set). B. The supernatants were concentrated using Amicon ultra filter device 3k MWCO. The flow through (MW <3k) and the concentrated sup (MW >3k) were used separately while inoculating WT biofilm. After 72 hours, crystal violet staining was performed to quantify the biofilm (Data are presented as mean ± ­SEM of 3 independent experiments). One-way ANOVA was used to analyze the data, p values *<0.05.

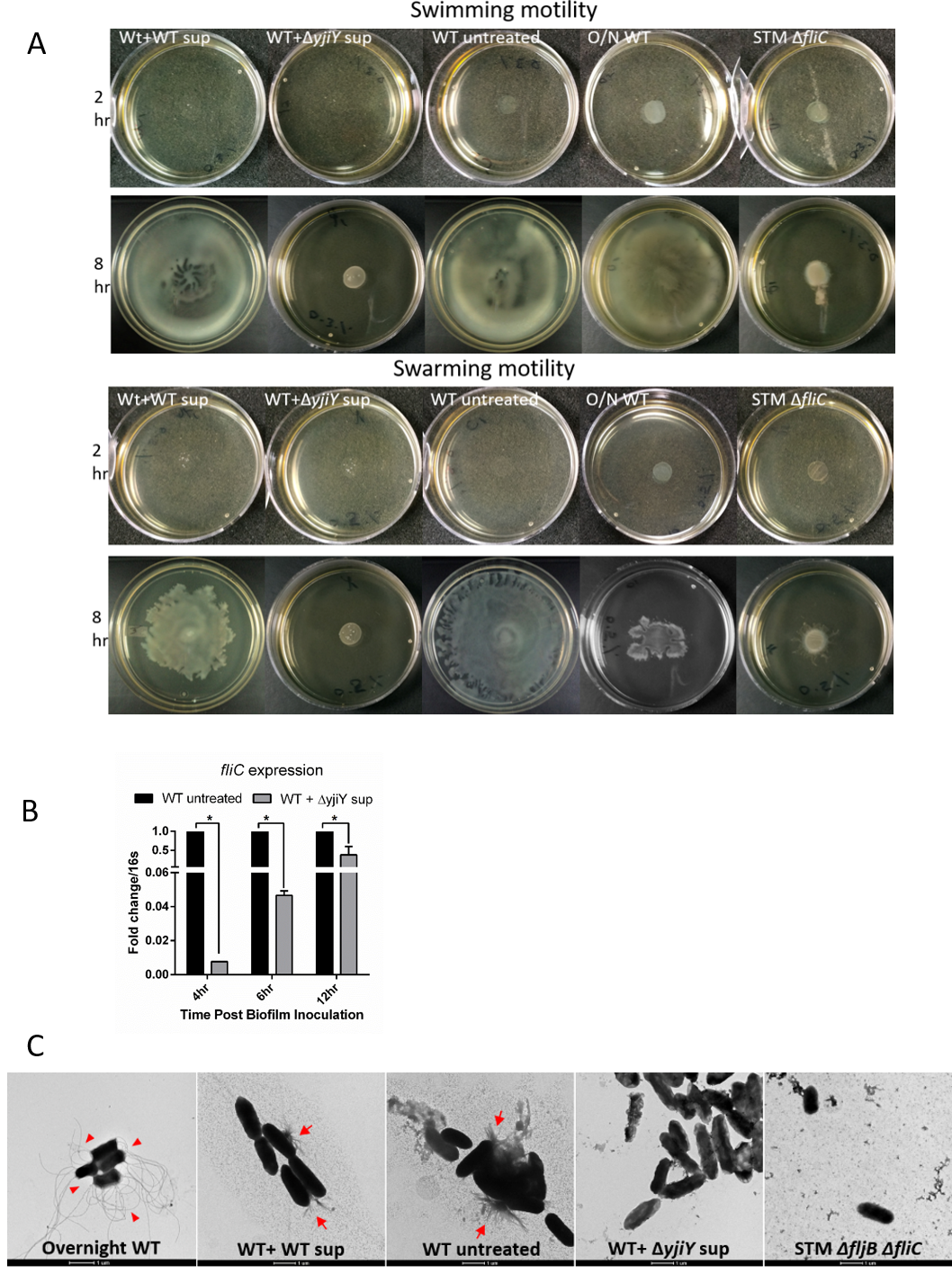

**Fig S4. Δ*yjiY* supernatant inhibits flagellar mediated bacterial motility**

A. Images of swimming and swarming plates inoculated with treated or untreated WT cells. Δ*fliC* after 2 hours and 8 hours post inoculation (Representative image from 3 independent experiments). B. *fliC* expression was checked from WT cells after 4 hours, 6 hours and 12 hours of inoculation with Δ*yjiY* supernatant in biofilm media (Data are presented as mean ± ­SEM of 2 independent experiments). Student’s t-test was used to analyze the data, p values *<0.05. C. Representative TEM images of STM WT cells inoculated with or without the supernatants. Cells were stained with uranyl acetate for visualization. Overnight grown STM WT cells and Δ*fljB* Δ*fliC* cultures were used positive and negative controls. Red arrowheads show intact flagella in overnight STM WT culture, red arrows show aggregates of fimbriae and fragmented flagella in untreated WT cells and WT cells treated with WT sup, after 72 hours of inoculation in biofilm media, whereas WT cells treated with Δ*yjiY* supernatant, do not show presence of such aggregates (Representative image from 2 independent experiments, approximately 50-60 cells were imaged from each experiment).

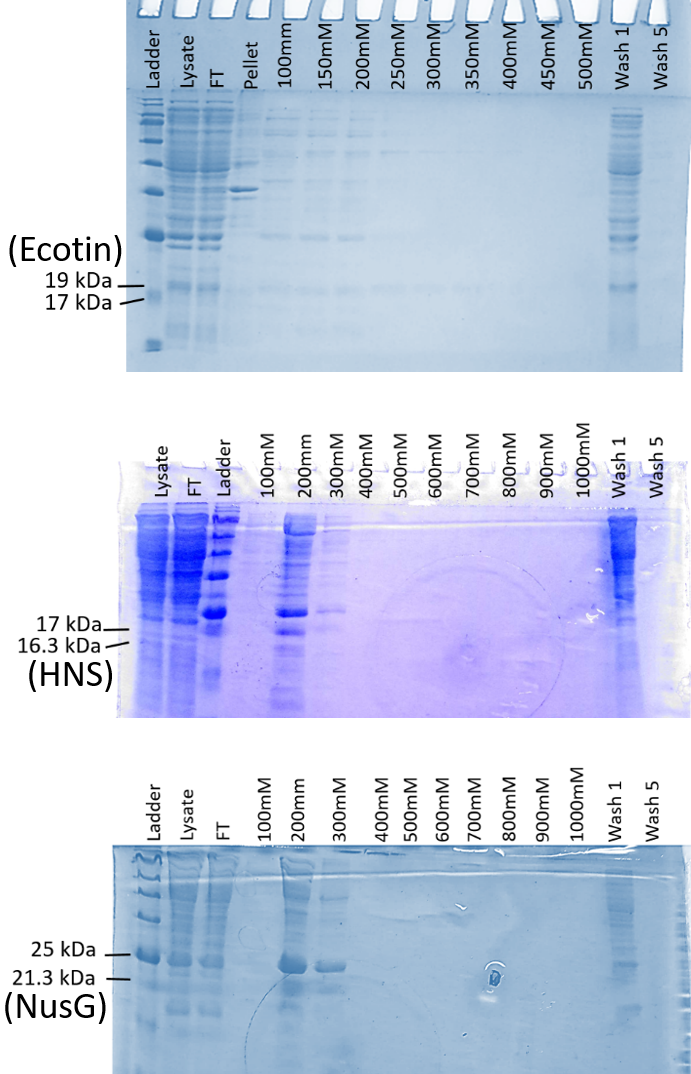

**Fig S5. Coomassie Brilliant Blue stained SDS-PAGE showing purified proteins of interest after elution from Ni-NTA affinity column with 100-1000mM imidazole**

*S.* Typhimurium 14028S Ecotin (STM14_2792), HNS (STM14_2116) and NusG (STM14_4985) genes were cloned with 6x His tag in pET15b vector using Gibson assembly. Positive clones were confirmed by PCR and transformed in *E. coli* BL21 (DE3) pLysS strain. After 1mM IPTG induction, cells were lysed by sonication on ice. The protein was purified from the soluble fraction of the lysate by affinity chromatography using Ni-NTA column. Protein was eluted with linear gradient of 100-1000mM imidazole in 50mM Tris. Top panel: Ecotin fractions (~19 kDa); middle panel: HNS fractions (~ 16.3 kDa); and bottom panel: NusG fractions (~21.3 kDa).

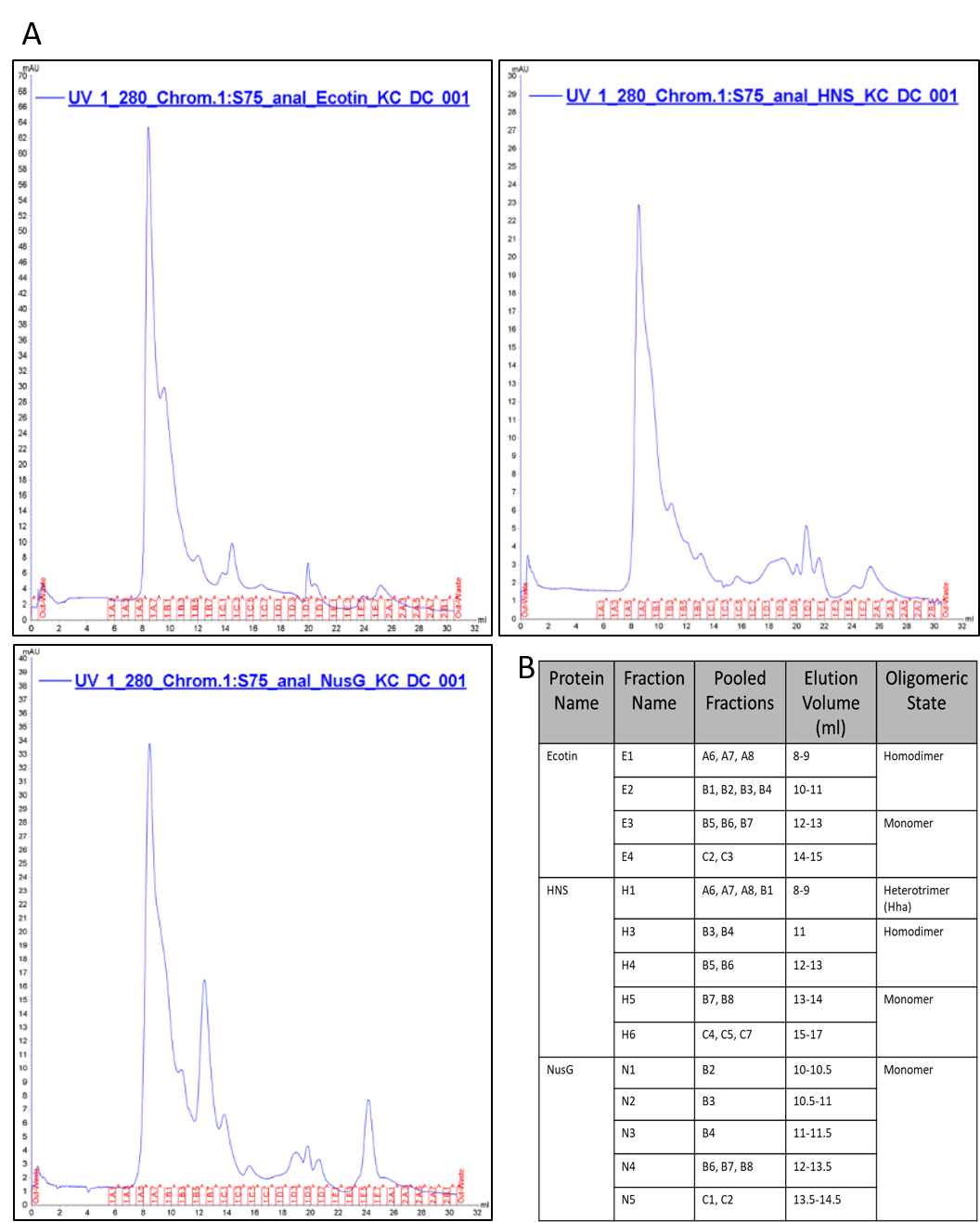

**Fig S6. Size exclusion chromatograms (SEC) of the pooled and concentrated fractions containing protein of interest**

A. Elute fractions containing protein of interest were pooled, concentrated and subjected to purification by size exclusion chromatography (SEC) on Superdex 75 Increase 10/300GL column (with a flow rate of 0.5 ml/min and 500 µl sample injection volume). B. Expected oligomeric state of proteins of interest (ecotin, HNS, and NusG) as predicted by ProtParam tool.

**Table S1.** List of proteins enriched in *ΔyjiY* supernatant.

| **UniProt ID** | **Proteins found in *yjiY* supernatant** | **Mol. Wt. (kDa)** |
| --- | --- | --- |
| tr\|A0A0F6AXD1 | Putative cytoplasmic protein | 11.8 |
| tr\|A0A0F6AZ15 | Putative ABC transporter periplasmic binding protein | 56.5 |
| tr\|A0A0F6AZ65 | Putative lipoprotein | 18.8 |
| tr\|A0A0F6AZA0 | Thioredoxin reductase | 34.8 |
| tr\|A0A0F6B006 | Anti-sigma28 factor **FlgM** | 10.5 |
| tr\|A0A0F6B054 | Transcription-repair-coupling factor **Mfd** | 129.9 |
| tr\|A0A0F6B0G8 | Putative ABC transporter periplasmic binding protein | 60.0 |
| tr\|A0A0F6B221 | Putative cytoplasmic protein | 18.6 |
| tr\|A0A0F6B2M8 | Probable transcriptional regulatory protein **YebC** | 26.4 |
| tr\|A0A0F6B3Z2 | Serine protease inhibitor **Ecotin** | 18.2 |
| tr\|A0A0F6B4D7 | Putative cytoplasmic protein | 10.2 |
| tr\|A0A0F6B9X9 | Transcription termination/antitermination protein **NusG** | 20.5 |
| tr\|A0A0F6BAJ5 | Putative outer membrane lipoprotein | 12.6 |
| tr\|A0A0F6BAL8 | Putative arginine-binding periplasmic protein | 27.4 |

**Table S2.** List of common proteins found in both the supernatants.

| **UniProt ID** | **Proteins found in both supernatants** | **Mol. Wt. (kDa)** | **Relative Abundance** (*ΔyjiY/*WT*)* |
| --- | --- | --- | --- |
| sp\|A0A0F6B244 | DNA-binding protein **H-NS** | 15.5 | 3.35 |
| tr\|A0A0F6AYA7 | Regulator of nucleoside diphosphate kinase **Rnk** | 14.99 | 3.30 |
| tr\|A0A0F6AYC2 | Cold shock protein **CspE** | 7.4 | 3.30 |
| tr\|A0A0F6B2E5 | Cold shock-like protein **CspC** | 7.4 | 5.48 |
| tr\|A0A0F6B5B6 | Heat shock protein/chaperone **GrpE** | 21.8 | 4.28 |
| tr\|A0A0F6B5K2 | DNA-binding protein **StpA** | 15.4 | 1.20 |
| tr\|A0A0F6B9S5 | ATP-dependent protease subunit **HslV** | 18.9 | 1.15 |
| tr\|A0A0F6AYJ2 | Flavodoxin **FldA** | 23.7 | 0.48 |
| tr\|A0A0F6B123 | Superoxide dismutase **SodB** | 21.3 | 0.72 |
| tr\|A0A0F6B125 | Glutaredoxin **YdhD** | 12.9 | 0.53 |
| tr\|A0A0F6B1W1 | Thiol peroxidase **Tph** | 18.0 | 0.37 |
| tr\|A0A0F6B2S0 | Ferritin **Ftn** | 19.3 | 0.10 |
| tr\|A0A0F6B4P5 | Thioredoxin-dependent thiol peroxidase **Bcp** | 17.6 | 0.80 |
| tr\|A0A0F6B7N9 | Bacterioferritin **Bfr** | 18.3 | 0.06 |
